# A longitudinal data resource to study brain development and transdiagnostic variation in executive function

**DOI:** 10.1101/2025.11.10.687633

**Authors:** Brooke L. Sevchik, Golia Shafiei, Kristin Murtha, Sophia Linguiti, Lia Brodrick, Juliette B.H. Brook, Matt Cieslak, Elizabeth Flook, Kahini Mehta, Steven L. Meisler, Kosha Ruparel, Sage Rush, Taylor Salo, S. Parker Singleton, Tien T. Tong, Mrugank Salunke, Dani S. Bassett, Monica E. Calkins, Mark A. Elliott, Raquel E. Gur, Ruben C. Gur, Tyler M. Moore, J. Cobb Scott, Russell T. Shinohara, M. Dylan Tisdall, Daniel H. Wolf, David R. Roalf, Theodore D. Satterthwaite

## Abstract

Executive function (EF) develops rapidly during adolescence. However, deficits in EF also emerge in adolescence, representing a transdiagnostic symptom associated with many forms of psychopathology. To promote transdiagnostic research on EF during development, we introduce a new data resource – the Penn Longitudinal Executive functioning in Adolescent Development study (Penn LEAD) – that combines longitudinal multimodal imaging data with rich clinical and cognitive phenotyping. These data include 225 imaging sessions from 132 individuals (8-16 years old at the time of enrollment) who are typically developing (27.3%), or meet criteria for attention-deficit hyperactivity disorder (20.5%) or the psychosis-spectrum (52.3%). In addition to phenotypic data from multiple cognitive tasks focused on EF, the study includes data from structural MRI, diffusion MRI, *n*-back task fMRI, resting-state fMRI, and arterial spin-labeled MRI. Notably, all raw data, fully-processed derived data, and detailed quality control recommendations are publicly shared on OpenNeuro. We anticipate that such analysis-ready data will accelerate research on EF development in psychiatry.

## Background & Summary/Introduction

Executive function (EF) is a broad cognitive domain, encompassing multiple component processes such as attentional control, working memory, abstraction, and set shifting. EF undergoes protracted development during adolescence and young adulthood as higher-order association cortex matures (Ferguson et al., 2021; Luna et al., 2010; Tervo-Clemmens et al., 2023). Deficits in EF often emerge in adolescence and are a major transdiagnostic domain of impairment across many forms of psychopathology (Powers & Casey, 2015; Romer & Pizzagalli, 2021; Shroff et al., 2024; Snyder et al., 2019). EF deficits notably play a central role in attention-deficit hyperactivity disorder (ADHD) and psychosis-spectrum (PS) disorders (Barkley, 1997; Forbes et al., 2009; Fusar-Poli et al., 2012). In these disorders and others, lower EF has been linked to greater rates of interpersonal conflict and poorer academic outcomes (De Wilde et al., 2016; Irwin et al., 2021; Jacobson et al., 2011; Morgan et al., 2019; Rohlf et al., 2018; Shanmugan et al., 2016). In this context, we introduce a new, open-access longitudinal data resource – Penn LEAD (Penn Longitudinal Executive functioning in Adolescent Development) – focused on EF in PS and ADHD.

This data resource is motivated by two gaps in the literature. First, while previous neuroimaging research in adults (Nowrangi et al., 2014) and children (Eng et al., 2022) has emphasized EF as a key marker of psychopathology, there are relatively few longitudinal neuroimaging data resources in adolescents with psychiatric disorders enriched for executive dysfunction. Many of the existing longitudinal datasets in adolescents focus mainly on healthy participants, excluding those with psychiatric disorders (developing Chinese Color Nest Project, Fan et al., 2023; Lifespan Human Connectome Project Development, Somerville et al., 2018; NIH-MRI of Normal Brain Development, Evans, 2006). The datasets that are focused on adolescent participants with psychiatric diagnoses often lack a wide breadth of multimodal imaging data, such as multi-shell diffusion-weighted imaging (DWI) and arterial spin-labeled MRI (ASL) (1000 Functional Connectomes Project, Biswal et al., 2010; ADHD-200 Consortium, 2012; Healthy Brain Network, Alexander et al., 2017; North American Prodrome Longitudinal Study, Addington et al., 2022; Oregon ADHD-1000, Nigg et al., 2023; Reproducible Brain Charts, Shafiei et al., 2025). Although the Adolescent Brain Cognitive Development Study (Karcher & Barch, 2021; Volkow et al., 2018), NKI-Rockland (Nooner et al., 2012; Tobe et al., 2022) and IMAGEN datasets (Schumann et al., 2010) provide rich multimodal data, they have relatively coarse measures of EF and were not specifically designed to assess EF development in the context of psychopathology.

A second gap that this resource seeks to address is the need for datasets that contain easily accessible, quality-controlled, analysis-ready data. Most datasets are restricted by cumbersome data-use agreements (DUAs) that can impede rapid progress (Shafiei et al., 2025; Di Martino et al., 2014; Jwa & Poldrack, 2022; Laird, 2021; Milham et al., 2018; Tedersoo et al., 2021; White et al., 2022; Zuo et al., 2014). Furthermore, many multimodal longitudinal imaging studies focused on adolescents with psychiatric disorders provide only raw or minimally-preprocessed data (Karcher & Barch, 2021; Schumann et al., 2010; Volkow et al., 2018). The lack of easily-accessible, fully-processed derivatives increases computational burden on researchers, leads to redundant efforts across research groups, and makes it difficult to reconcile disparate findings across studies.

To address these gaps, Penn LEAD combines longitudinal, fully-processed, multimodal imaging data with extensive cognitive and clinical phenotyping focused on EF. Multimodal neuroimaging data includes structural magnetic resonance imaging (sMRI), diffusion-weighted imaging (DWI), functional MRI (fMRI; resting-state and *n*-back task), ASL, and a multi-echo gradient-recalled echo sequence (MEGRE) for quantitative susceptibility mapping (QSM). In addition to raw data, this data resource includes fully-processed, derived data from open-source workflows as well as detailed assessments of image quality. To maximize reproducibility, all processed data were generated using the “FAIRly-big” framework (Wagner et al., 2022), which provides full provenance-tracking with DataLad (Halchenko et al., 2021). Anonymized data are openly available on OpenNeuro without a DUA, which removes barriers to data access and facilitates replication of analyses between different research groups. As described below, we anticipate that this dataset will accelerate research on transdiagnostic variation of EF in adolescence.

## Methods

### Recruitment, Participants, and Procedures

Participants were identified and enrolled in the study across two study visits (see **Figure 1** for study timeline). First, participants were recruited from the Children’s Hospital of Philadelphia (CHOP) patient population, via medical record review and referrals from outpatient behavioral health or primary care clinics. Our target population included youth between 8-16 years old. Exclusion criteria consisted of: 1) metallic implants, claustrophobia, or other contraindications to MRI; 2) significant medical or neurological illness that impacted brain function or impeded participation; 3) acute intoxication with alcohol or other substances based on clinical assessment or participant report; and 4) pregnancy. Participant demographics are detailed in **Figure 2c**. Participants were invited to participate in a two-stage research study involving neuroimaging as well as clinical and cognitive phenotyping.

**Figure 1.**
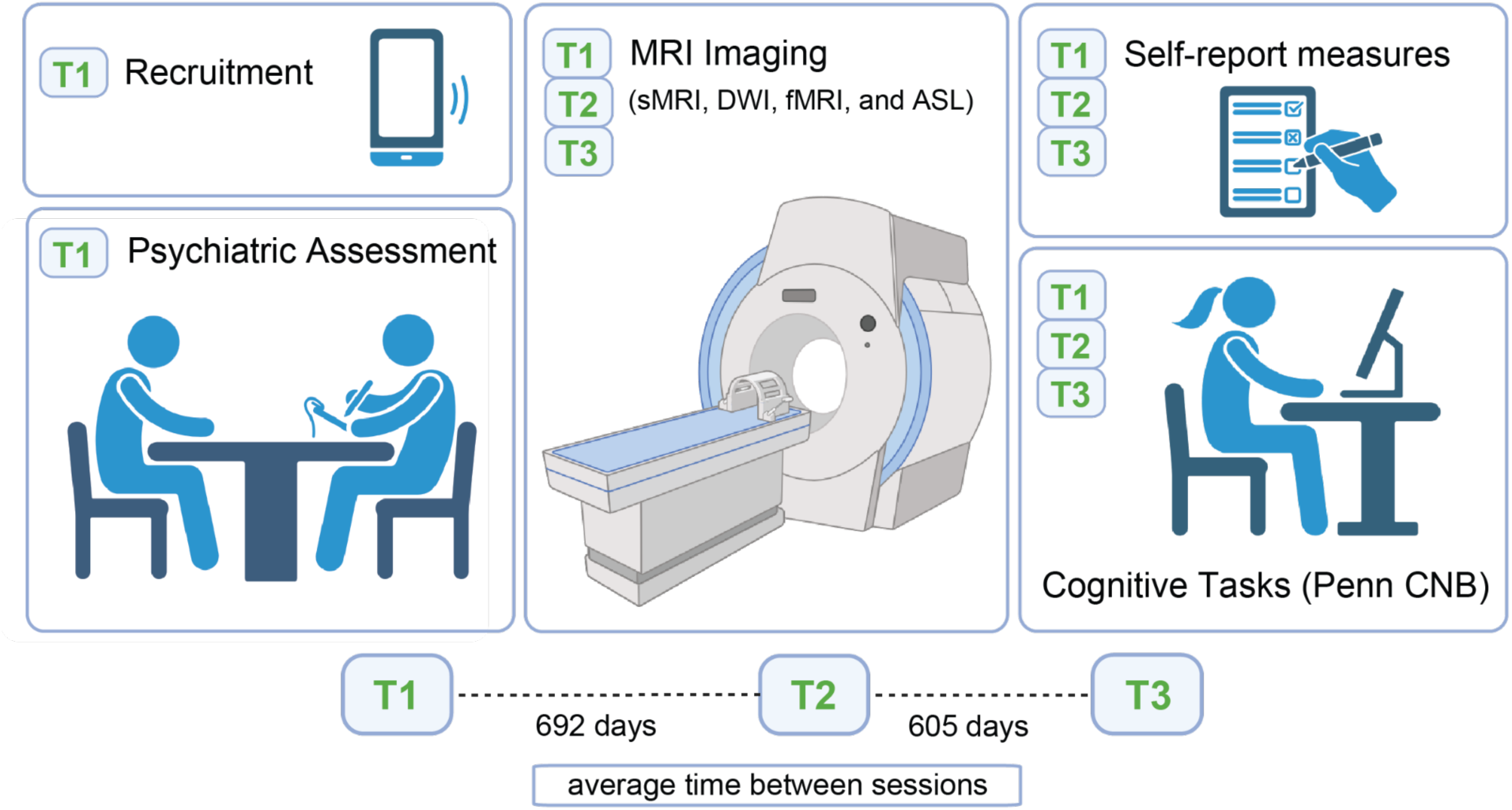
Overview of recruitment and visits. At Timepoint 1 (T1), participants were recruited from the Children’s Hospital of Philadelphia, then were placed into ‘ADHD,’ ‘Psychosis Spectrum (PS),’ or ‘Typically Developing’ study groups following psychiatric assessment via a detailed clinical interview. At T1, in addition to any subsequent longitudinal timepoints (T2 and T3), participants completed magnetic resonance imaging (MRI) scanning sessions, self-report assessments, and the Penn Computerized Neurocognitive Battery (Penn CNB). sMRI = structural MRI; DWI = diffusion-weighted MRI; fMRI = functional MRI; ASL = arterial spin labeling. Created in BioRender. Sevchik, B. (2025) https://BioRender.com/ern7vq6

**Figure 2.**
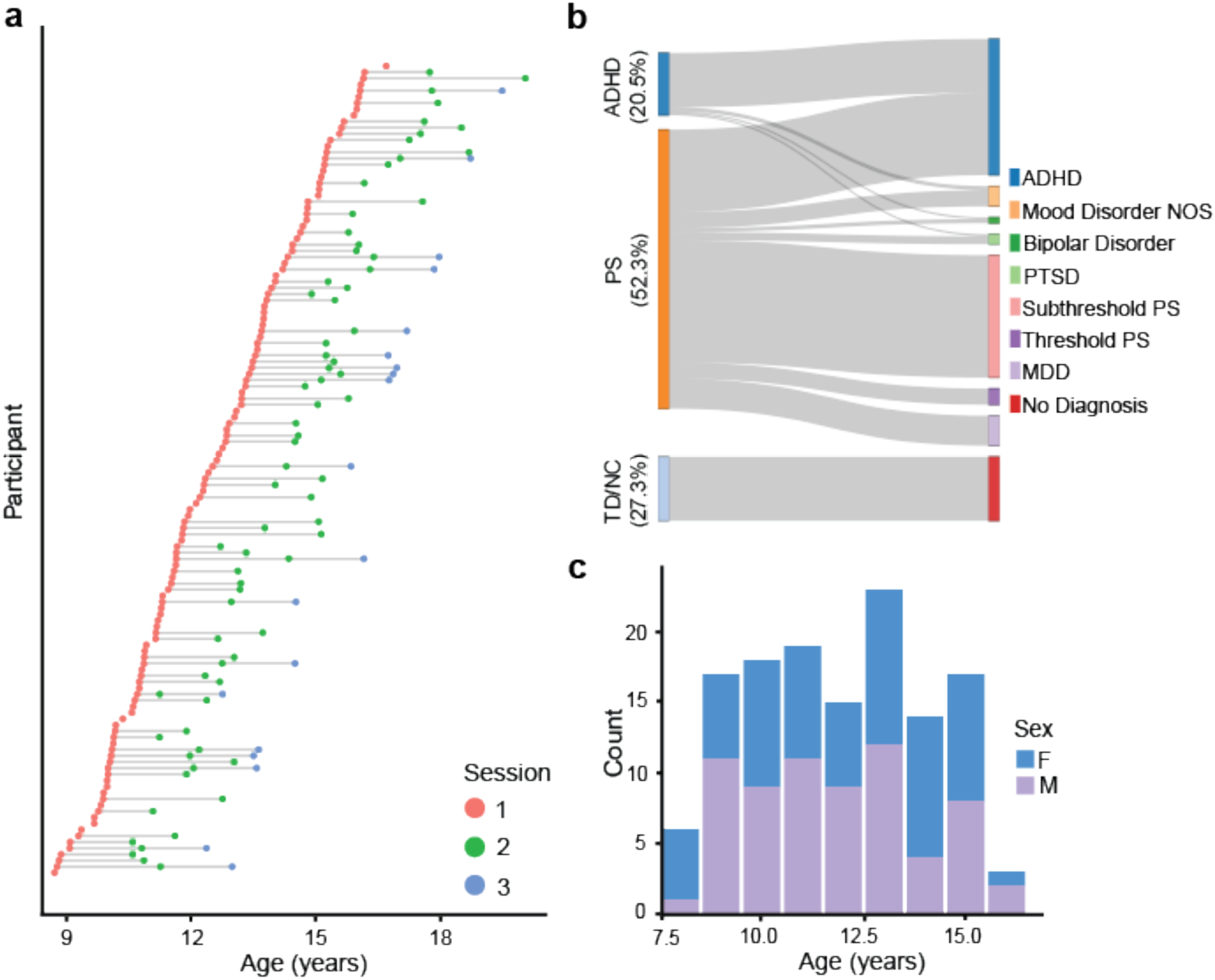
Participant demographics and clinical diagnostic summary. **(a)** Longitudinal data is represented as a feather plot, where each individual dot represents an individual session. Line lengths between sessions are proportional to the time elapsed between sessions for each participant. **(b)** Clinical diagnostic summary data are visualized using a sankey plot. Values on the left side of the sankey plot correspond to the percentage of unique participants in each study group out of all participants in the dataset, according to clinical interviews (ADHD = attention-deficit hyperactivity disorder; PS = psychosis-spectrum (including threshold and subthreshold); TD = typically developing). Rectangles on the right of the plot correspond to the percentage of unique participants who met diagnostic criteria for each disorder out of the total sample: ADHD: 51.5%; Mood Disorder Not Otherwise Specified (Mood Disorder NOS): 7.6%; Bipolar disorder: 2.3%; Post-traumatic stress disorder (PTSD): 3.8%; Subthreshold Psychosis Spectrum (PS; clinical high risk for psychosis): 46.2%; Threshold Psychosis Spectrum (PS): 6.1%; Major depressive disorder (MDD): 11.4%; No clinical psychiatric diagnosis (No Diagnosis): 27.3%. Gray lines from the left to the right of the sankey plot are proportional to the amount of participants from each study group contributing to the total percentage of participants with each disorder. **(c)** The distribution of the age at intake and sex of the sample: Female: 49.2%; Average age at intake: 11.9 years old.

During the first visit, informed consent for study procedures was provided by the parent or guardian of minor participants; minor participants provided assent. The informed consent document contained language from the Open Brain Consent (Halchenko, 2022) to facilitate open data sharing. Study procedures were approved by the University of Pennsylvania and Children’s Hospital of Philadelphia Institutional Review Boards (application number 829744).

Subsequently, youth and their parents completed an initial 3-5 hour clinical visit during which participants received a detailed diagnostic assessment from trained assessors (Calkins et al., 2015) (**Figure 2b**). Parents provided collateral information for all participants; children over the age of eleven years were also interviewed about their own symptoms. The computerized assessment combined a modified and adapted version of Kiddie Schedule for Affective Disorders and Schizophrenia (K-SADS; Kaufman et al., 1997; see Calkins et al., 2017 for details on the modifications) as well as the Structured Interview for Prodromal Symptoms (SIPS; Miller et al., 2003). During this first visit, participants also completed a broad battery of cognitive tasks (see **Cognitive Assessment**, below; Gur et al., 2010, 2012; Moore et al., 2015, 2023; Scott et al., 2020). Diagnostic and neurocognitive testing information was collected from *N* = 132 participants (49.2% female; average age at intake = 11.9 years).

On the second visit, youth were invited to participate in neuroimaging procedures if the semi-structured diagnostic assessment during the first visit indicated that they met criteria for one of three diagnostic groups. Specifically, youth were asked to participate in neuroimaging if they met criteria for ADHD, PS, or were typically-developing (TD, i.e., no current or lifetime history of any DSM-V diagnosis). Co-morbidity was permitted; participants meeting criteria for both PS and ADHD diagnoses were considered in the psychosis-spectrum study group for enrollment purposes.

Participants completed a 4-5 hour neurocognitive and neuroimaging visit at each longitudinal timepoint, with one to three neuroimaging visits per participant (average time between sessions = 648.5 days; **Figure 2a**). At each neuroimaging visit participants completed cognitive testing focused on executive functioning tasks (see **Cognitive Assessment**) and a battery of self-report surveys (see **Self-Report Data**). Participants with a second and/or third neuroimaging visit repeated the same battery of cognitive tests as during their first visit. In total, this dataset includes 225 neuroimaging sessions from *N* = 132 participants (49.2% female, average age across all scans = 13.4 years).

### Cognitive Assessment

Cognition was assessed with the Penn Computerized Neurocognitive Battery (Penn CNB; Gur et al., 2010, 2012; Moore et al., 2015, 2023; Scott et al., 2020). In this study, we included both tasks measuring general cognition (Lifespan Brain Institute CNB; LiBI CNB) and a specialized suite of tasks focused on executive function ability (EF CNB). In total, these two batteries comprised 21 distinct tasks (**Table 2)**. For timepoint 1, the LiBI CNB was administered on the day of the clinical interview, and the EF CNB was administered on the day of the MRI scan. For longitudinal timepoints 2 and 3, both the LiBI CNB and the EF CNB were administered on the day of the MRI scan.

**Table 1.** Neuroimaging acquisition parameters.

| <u>Sequence</u> | <u>Slices</u> | <u>%</u><br><u>FOV</u><br><u>phase</u> | <u>Resolution</u><br><u>(mm)</u> | <u>TR</u><br><u>(ms)</u> | <u>TE</u><br><u>(ms)</u> | <u>TI</u><br><u>(ms)</u> | <u>Flip</u><br><u>Angle</u><br><u>(degree)</u> | <u>Multi</u><br><u>Band</u><br><u>Accel</u> | <u>Phase</u><br><u>Partial</u><br><u>Fourier</u> |
| --- | --- | --- | --- | --- | --- | --- | --- | --- | --- |
| T1w | 176 | 100% | 1.0x1.0x1.0 | 2500 | 2.9 | 1070 | 8 | n/a | 1.0 |
| T2w | 176 | 100% | 1.0x1.0x1.0 | 3200 | 565 | n/a | 120 | n/a | 1.0 |
| DWI | 81 | 100% | 1.7x1.7x1.7 | 4200 | 89 | n/a | 90 | 3 | 0.75 |
| DWI<br>fmap PA | 81 | 100% | 1.7x1.7x1.7 | 12400 | 89 | n/a | 90 | n/a | 0.75 |
| DWI<br>fmap AP | 81 | 100% | 1.7x1.7x1.7 | 12400 | 89 | n/a | 90 | n/a | 0.75 |
| Resting-state<br>fMRI | 60 | 100% | 2.4x2.4x2.4 | 800 | 30 | n/a | 52 | 6 | 1.0 |
| *n*-back<br>task fMRI | 60 | 100% | 2.4x2.4x2.4 | 800 | 30 | n/a | 52 | 6 | 1.0 |
| fMRI<br>fmap PA | 60 | 100% | 2.4x2.4x2.4 | 7030 | 80 | n/a | 90 | n/a | 1.0 |
| fMRI<br>fmap AP | 60 | 100% | 2.4x2.4x2.4 | 7030 | 80 | n/a | 90 | n/a | 1.0 |
| ASL | 32 | 100% | 3.75x3.75x3.75 | 4298 | 10 | n/a | 90 | n/a | 1.0 |
| MEGRE | 96 | 100% | 1.5x1.5x1.5 | 35 | 7.5,<br>15,<br>22.5,<br>30 | n/a | 15 | n/a | 1.0 |

**Table 2.** Overview of cognitive tasks in the dataset from the Penn CNB. The table above summarizes the cognitive domains corresponding to the tasks included in batteries from the Penn Computerized Neurocognitive Battery that we included in our dataset. Detailed descriptions of each task can be found in the papers cited for each task.

| **CNB Task** | **Battery** | **Cognitive Domain** | **Citation** |
| --- | --- | --- | --- |
| Penn Abstraction, Inhibition and Working Memory Task | EF | Executive function | Moore et al., 2023; Glahn et al., 2000 |
| Penn Face Memory Test | LiBI | Episodic memory | Moore et al., 2015, 2023; Gur et al., 2010, 2012; Scott et al., 2020 |
| Children's Penn Word Memory Test | LiBI | Episodic memory | Moore et al., 2015, 2023; Gur et al., 2010, 2012 |
| Short Computerized Finger-Tapping Task | LiBI | Sensorimotor speed | Gur et al., 2010, 2012; Scott et al., 2020 |
| Penn Emotion Recognition Task for Children | LiBI | Social cognition | Moore et al., 2015, 2023; Gur et al., 2010, 2012 |
| Digit Symbol Test | EF | Executive function | Scott et al., 2020; Moore et al., 2023 |
| Short Letter-*N*-Back (2-back) | LiBI | Executive function | Moore et al., 2015; Gur et al., 2010; Gur et al., 2012 |
| Measured Emotion Differentiation Test | LiBI | Social cognition | Moore et al., 2015; Gur et al., 2012; Moore et al., 2023 |
| Motor Praxis Test | LiBI | Sensorimotor speed | Gur et al., 2012; Scott et al., 2020 |
| Penn Conditional Exclusion Task for Children | LiBI | Executive function | Moore et al., 2015; Gur et al., 2010, 2012; Scott et al., 2020 |
| Short Penn Continuous Performance Test-Number and Letter Version | LiBI | Executive function | Moore et al., 2015, 2023; Gur et al., 2010, 2012; Scott et al., 2020 |
| Penn Matrix Reasoning Test | LiBI | Complex cognition | Moore et al., 2015, 2023; Gur et al., 2012; Scott et al., 2020 |
| Variable Short Penn Line Orientation Test | LiBI | Complex cognition | Moore et al., 2015, 2023; Gur et al., 2010, 2012; Scott et al., 2020 |
| Children's Verbal Reasoning Test | LiBI | Complex cognition | Moore et al., 2015, 2023; Gur et al., 2010, 2012 |
| Short Visual Object Learning Test | LiBI | Episodic memory | Moore et al., 2015, 2023; Gur et al., 2010, 2012; Scott et al., 2020 |
| Trailmaking A | EF | Executive function | Scott et al., 2020 |
| Trailmaking B | EF | Executive function | Scott et al., 2020 |
| Effort Discounting Task | EF | Reward | Moore et al., 2023 |
| Delayed Discounting Task | EF | Reward | Moore et al., 2023 |
| Risk Discounting Task | EF | Reward | Moore et al., 2023 |
| Age Differentiation Test | LiBI | Social cognition | Moore et al., 2015, 2023; Gur et al., 2012 |

The CNB includes summary measures of cognitive performance based on accuracy and response time. These include a rich variety of measures, such as the median response times as well as the sum (and/or average) of correct responses, incorrect responses, true positives, false positives, false negatives, and true negatives. The data provided are not normalized. Detailed descriptions of each task can be found in the citations in **Table 2**.

### Self-Report Data

All participants completed a broad array of self-report batteries (**Table 3**). Scales were grouped into main, pre-scan, and post-scan batteries. The main battery consisted of the Adverse Life Experiences Scale (ALES; Hawes et al., 2021), Affective Lability Scale (ALS-18; Contardi et al., 2018), Affective Reactivity Index (ARI; Stringaris et al., 2012), Altman Self-Rating Mania Scale (ASRM; Altman et al., 1997), Beck Depression Inventory (BDI-1A; Beck, 1961, modified child version without suicide item), Behavioral Inhibition/Activation System (BIS-BAS child version; Carver & White, 1994; Muris et al., 2005), the Extended Strengths and Weaknesses Assessment of Attention-Deficit/Hyperactivity-Disorder Symptoms and Normal Behavior (E-SWAN ADHD; Alexander et al., 2019; Swanson et al., 2012), the Extended Strengths and Weaknesses Assessment of Disruptive Mood Dysregulation Disorder Symptoms and Normal Behavior (E-SWAN DMDD; Alexander et al., 2020), The Motivation and Pleasure Scale - Self Report (MAP-SR; Llerena et al., 2013), Positive and Negative Affect Schedule (PANAS child; Watson et al., 1988), PROMIS Pediatric Anxiety scale (PPA; Irwin et al., 2012), PRIME Screen (self-report) (Calkins et al., 2025; Kobayashi et al., 2008), Revised Physical Anhedonia Scale (RPAS; Chapman et al., 1976), Revised Social Anhedonia Scale (RSAS; Chapman et al., 1976), Substance Use Questionnaire (Han et al., 1999), Tanner Developmental Scale (Boy) (Marshall & Tanner, 1970), Tanner Developmental Scale (Girl) (Marshall & Tanner, 1969), and Wolf Intrinsic Motivation/Extrinsic Motivation Scale (Wolf IM/EM short version; Didier et al., 2023). Descriptions of each scale can be found in the citations in **Table 3**. Participants completed the pre-scan battery before their MRI scan, consisting of the full State-Trait Anxiety Inventory (STAI; Spielberger et al., 1983), and the post-scan battery after their MRI scan, consisting of only the state items (items 1-20) of the STAI after their MRI scan.

**Table 3.** Overview of self-report scales in the dataset. The table above summarizes the self-report scales collected, along with the symptom or trait they are designed to measure. The full STAI was administered before the scan as part of the pre-scan battery, and just the state anxiety items of the STAI were administered after the scan as part of the post-scan battery. All other scales were administered as part of the main battery at each session. Details about each scale can be found in the papers cited for each scale

| **Self-Report Scale** | **Battery** | **Measures** | **Citation** |
| --- | --- | --- | --- |
| Adverse Life Experiences Scale (ALES) | main | Occurrence and developmental timing of adverse childhood experiences | Hawes et al., 2021 |
| Affective Lability Scale (ALS-18) | main | Affective lability (aspect of emotion dysregulation) | Contardi et al., 2018 |
| Affective Reactivity Index (ARI) | main | Irritability | Stringaris et al., 2012 |
| Altman Self-Rating Mania Scale (ASRM) | main | Mania | Altman et al., 1997 |
| Beck Depression Inventory (BDI-1A) | main | Depression | Beck, 1961 |
| Behavioral Inhibition/Activation System (BIS-BAS) | main | Motivational systems underlying behavior and affect | Carver & White, 1994 |
| Extended Strengths and Weaknesses Assessment of ADHD Symptoms and Normal Behavior (E-SWAN ADHD) | main | ADHD symptoms | Alexander et al., 2020; Swanson et al., 2012 |
| Extended Strengths and Weaknesses Assessment of DMDD Symptoms and Normal Behavior - DMDD (E-SWAN DMDD) | main | DMDD symptoms | Alexander et al., 2020 |
| The Motivation and Pleasure Scale - Self-Report (MAP-SR) | main | Severity of negative symptoms | Llerena et al., 2013 |
| Positive and Negative Affect Schedule (PANAS) | main | Positive and negative affect (mood) | Watson et al., 1988 |
| PROMIS Pediatric Anxiety (PPA) | main | Anxiety symptoms | Irwin et al., 2012 |
| PRIME Screen (Self-Report) | main | Subthreshold psychosis symptoms | Kobayashi et al., 2008, Calkins et al., 2025 |
| Revised Physical Anhedonia Scale (RPAS) | main | Physical anhedonia symptoms | Chapman et al., 1976 |
| Revised Social Anhedonia Scale (RSAS) | main | Social anhedonia symptoms | Chapman et al., 1976 |
| Substance Use Questionnaire | main | Substance use | Han et al., 1999 |
| Tanner Developmental Scale - Boy | main | Pubertal changes in boys | Marshall & Tanner, 1970 |
| Tanner Developmental Scale - Girl | main | Pubertal changes in girls | Marshall & Tanner, 1969 |
| Wolf Intrinsic Motivation/Extrinsic Motivation (Wolf IM/EM) | main | Trait intrinsic and extrinsic motivation | Didier et al., 2023 |
| State-Trait Anxiety Inventory (state) | pre-scan<br>post-scan | State anxiety | Spielberger et al., 1983 |
| State-Trait Anxiety Inventory (trait) | pre-scan | Trait anxiety | Spielberger et al., 1983 |

We release the raw self-report data with their summary scores (where applicable) for each session. We did not include summary scores for the Tanner Developmental Stages scales due to complexities in how puberty is staged, and we did not include summary scores for the Substance Use Questionnaire given the lack of participants who endorsed drug use and the complexity of substance use. Summary scores were calculated in accordance with guidelines from each scale’s original publication, primarily by calculating the sums and averages of individual item-level data. For each scale, we did not calculate a summary score for a given session if it was missing any individual item. Scoring details are described in the JSON files accompanying each scale. The code used for scoring is also available publicly on GitHub.

### Neuroimaging Data Acquisition

MRI data were acquired on a 3T Siemens Prisma scanner (Erlangen, Germany) with a 32-channel head coil at the University of Pennsylvania. The imaging protocol was approximately 60 minutes long. The full MRI protocol is provided as a PDF file on OpenNeuro, alongside the complete dataset. Prior to scanning, participants completed a brief practice version of the *n*-back task to prepare them for performing the same task in the MRI scanner.

During the scan, participants were placed supine in the scanner and were given earplugs and head padding to both muffle noise and reduce head motion. Sequences included sMRI T1-weighted (T1w) and T2-weighted (T2w) scans; fMRI resting-state and *n*-back task-based scans; ASL scans; DWI scans; and a MEGRE sequence for QSM. Raw imaging data includes a total of 223 T1w scans, 224 T2w scans, 213 resting-state fMRI scans, 209 task fMRI scans, 188 ASL scans, 219 diffusion scans, and 181 MEGRE scans (**Figure 4**). A summary of sequence parameters is provided in **Table 1**.

#### Structural MRI

We first acquired T1w structural images using acquisition parameters that were highly similar to the ABCD study (Karcher & Barch, 2021; Volkow et al., 2018: 176 slices; repetition time, TR = 2500 ms; TI = 1070 ms; echo time, TE = 2.9 ms; flip angle, FA = 8°; field of view, FOV = 256 x 256 mm; matrix size = 256 x 256; voxel size = 1.0 mm; phase encoding direction = A >> P; acquisition time 7:12 min). Similarly, the T2w structural scan was harmonized with the ABCD study (176 slices; repetition time, TR = 3200 ms; echo time, TE = 565 ms; flip angle, FA = 120°; field of view, FOV = 256 x 256 mm; matrix size = 256 x 256; voxel size = 1.0 mm; phase encoding direction = A >> P; acquisition time 6:35 min). During structural scans, participants watched movie clips from *Harry Potter* or *Ratatouille*, depending on their preference.

#### Diffusion MRI

We also acquired a multi-shell, single spin-echo DWI sequence in the Posterior-to-Anterior phase encoding direction, using a monopolar sampling scheme, with acquisition parameters highly similar to the ABCD study (Karcher & Barch, 2021; Volkow et al., 2018: repetition time, TR = 4200 ms; echo time, TE = 89 ms; field of view, FOV = 240 x 240 mm; matrix = 140 x 140; voxel size = isotropic 1.7 mm; 81 interleaved slices acquired posterior to anterior; a multiband factor of 3; acquisition time = 7:29 min). We acquired the following volumes: 7 at *b* = 0, 6 at *b* =500, 15 at *b* =1000, 15 at *b* =2000, and 60 at *b* = 3000 s/mm^2^ (103 volumes total). These diffusion sequences were accompanied by an additional *b* = 0 scan acquired in both the Posterior-to-Anterior and Anterior-to-Posterior phase encoding directions (repetition time, TR = 12400 ms; echo time, TE = 89 ms; flip angle, FA = 90°; field of view, FOV = 240 x 240 mm; voxel size = isotropic 1.7 mm; 81 interleaved slices; multiband factor = 1; acquisition time = 0:25 min each) that were used to estimate a field map.

#### Functional MRI

Acquisition parameters for fMRI scans were also highly similar to the ABCD study (Karcher & Barch, 2021; Volkow et al., 2018). We acquired one run of *n*-back task fMRI, two runs of resting-state fMRI, and accompanying field maps. During the fMRI *n*-back task run (single-echo; 60 slices; repetition time, TR = 800 ms; echo time, TE = 30 ms; flip angle, FA = 52°; field of view, FOV = 216 x 216 mm; matrix size = 90 x 90; voxel size = isotropic 2.4 mm; multiband factor = 6; in-plane acceleration factor = 1; 522 volumes; acquisition time = 7:02 min), participants completed a version of the *n*-back task with fractal images (Ragland et al., 2015; Satterthwaite et al., 2013b). In the scanner, participants were shown a continuous sequence of fractal images and were instructed to press a button if the current image matched the image they saw *n* stimuli before. We used a modified version of the *n*-back task, during which participants completed the task for two working memory levels: 0-back, for which they pressed the button when the fractal image matched a specified target fractal image; and 2-back, for which they pressed the button if the fractal image matched the image from two trials before it. We did not include an intermediate 1-back task. Scored task event files can be found with the raw fMRI data on OpenNeuro.

We also acquired two runs of fMRI where participants were asked to view a fixation cross (60 slices; repetition time, TR = 800 ms; echo time, TE = 30 ms; FA=52°; FOV = 216 mm; matrix size = 90 x 90; voxel size = isotropic 2.4 mm; multiband factor = 6; in-plane acceleration factor = 1). The first run (522 volumes, acquisition time = 7:02 min) was acquired before the fractal *n*-back task run; the second run (383 volumes, acquisition time = 5:11 min) was acquired after the fractal *n*-back task. In addition, field maps with one volume in both the Anterior-to-Posterior and Posterior-to-Anterior phase encoding direction were collected for distortion correction (60 slices; repetition time, TR = 7030 ms; echo time, TE = 80 ms; flip angle, FA = 90°; field of view, FOV = 216 x 216 mm; matrix size = 90 x 90; voxel size = isotropic 2.4 mm; acquisition time 0:14 min each).

#### Arterial spin-labeled MRI

We measured cerebral perfusion using an unbalanced, background-suppressed, 3D stack of spirals pseudo-continuous arterial spin-labeled (PCASL) MRI scan (labeling duration = 1500 ms; post-labeling delay, PLD = 1500 ms; repetition time, TR = 4298 ms; echo time, TE = 10 ms; field of view, FOV = 240 x 240 mm; voxel size = isotropic 3.8 mm; 80 volumes; acquisition time = 5:43 min). This acquisition was accompanied by a separate M0 reference scan with the same spatial parameters (repetition time, TR = 4000 ms; echo time, TE = 10.03 ms).

#### MEGRE

We also acquired a MEGRE sequence with four echos (96 slices; repetition time, TR = 35 ms; echo times, TEs = 7.50, 15.00, 22.50, 30.00 ms; flip angle, FA = 15°; field of view, FOV

= 240 x 180 mm; matrix size = 160 x 120; voxel size = isotropic 1.5 mm; phase encoding direction = R >> L; acquisition time 3:59 min). This scan can be used for QSM.

### Neuroimaging Curation

We used CuBIDS software (Curation of BIDS; Covitz et al., 2022) to curate the raw neuroimaging data and metadata. CuBIDS allows for fully reproducible curation of metadata and file names in the BIDS dataset using DataLad (Halchenko et al., 2021). Notably, BIDS Apps used for image processing are automatically configured based on the metadata they encounter. As such, if metadata are incorrect, preprocessing pipelines run the risk of being “reproducible but wrong.” Accordingly, careful metadata curation with CuBIDS helps ensure accurate preprocessing pipelines, thereby reducing the potential for inaccurate preprocessing pipelines due to absent or incorrect metadata. All code for metadata curation is available on GitHub.

CuBIDS also helps identify sources of variation and heterogeneity in the imaging data. For each scan type, the ‘Dominant Group’ represents the subset of acquisition parameters that match the majority of files, whereas ‘Variant Groups’ denote the subset of scans with different acquisition parameters (**Table S1)**. For example, among *n*-back scans, *N* = 1 had a lower number of volumes than expected, and was assigned to a Variant Group. In contrast, the majority (*N* = 185) were acquired with the expected number of volumes with no other metadata variation, and assigned to the Dominant Group. The full CuBIDS summary file is available on OpenNeuro (ds006688, `code/` subfolder, v50). ASL scans have a lower percentage of scans (60.11%) in the dominant group; this is due to minor variances in effective echo spacing, repetition time, and total readout time. Most scans not in the dominant group for fMRI runs (11.5% for *n-*back, 12.1% for the first resting state run, 15.6% for the second resting state run) are due to small amounts of variation in scan obliquity or the number of volumes acquired.

### Image Processing

As data are openly shared without a DUA, maintaining participant data privacy is of paramount concern. Accordingly, we anonymized structural images using AFNI’s refacer (Cox, 1996) for the T1w images and Pydeface v2.0.2 (Gulban et al., 2022) for the T2w images. Following anonymization, we utilized BIDS Apps (Gorgolewski et al., 2016) for image processing including sMRIPrep (Esteban et al., 2018, 2019, & 2020), fMRIPrep (Esteban et al., 2018, 2019, & 2020), QSIPrep (Cieslak et al., 2021), QSIRecon (Cieslak et al., 2021), and ASLPrep (Adebimpe et al., 2022, 2023). All BIDS Apps for image processing were run via the BIDS-App Bootstrap software (BABS; Zhao et al., 2024) that uses DataLad (Halchenko et al., 2021) to provide a full audit trail and enhance reproducibility. We utilized the University of Pennsylvania’s CUBIC computing resource, a RedHatEnterprise Linux-based HPC cluster. Note that we did not complete any processing of the MEGRE scans; the raw data is available on OpenNeuro. **Figure 4** describes the number of scans available at each stage, including raw data, processed images, and scans passing quality control (QC; see **Quality Control**). Detailed information on image processing compiled below are derived from methods descriptions generated by each BIDS App, under the CC0 1.0 Universal license.

#### Structural image processing

Anatomical images were processed using sMRIPrep, executed as the anatomical-only option in fMRIPrep v25.0.0 (Esteban et al., 2018, 2019, & 2020) based on Nipype v1.9.2 (Gorgolewski et al., 2011 & 2018). Using the sMRIPrep workflow allowed us to use the anatomical outputs in multiple downstream workflows, such as fMRIPrep and ASLPrep. As part of sMRIPrep, the T1w image was corrected for intensity non-uniformity (INU) with N4BiasFieldCorrection (Tustison et al., 2010) with ANTs 2.5.4 (Avants et al., 2008). The image was then skull-stripped with a Nipype implementation of the antsBrainExtraction.sh workflow. Brain tissue segmentation was performed on the brain-extracted T1w using fast (FSL v6.0.7.7; Zhang, Brady, and Smith 2001). Brain surfaces were reconstructed using FreeSurfer’s recon-all workflow (FreeSurfer 7.3.2; Dale, Fischl, and Sereno, 1999). The brain mask estimated previously with the ANTs brain extraction was refined to reconcile ANTs-derived and FreeSurfer-derived segmentations. The T2w image was used to improve pial surface refinement. Volume-based spatial normalization to two standard spaces (MNI152NLin6Asym, MNI152NLin2009cAsym) was performed using antsRegistration (ANTs 2.5.4), using brain-extracted versions of both T1w reference and the T1w template. Grayordinate “dscalar” files containing 91k grayordinates were resampled onto fsLR 32K space using Connectome Workbench (Glasser et al., 2013). Outputs from sMRIPrep were further processed using freesurfer-post v0.1.2 (https://github.com/PennLINC/freesurfer-post.git). Freesurfer-post parcellates and then aggregates morphometric measures from FreeSurfer into tabulated data to facilitate cross-modality comparisons. FreeSurfer data was parcellated using multiple common atlases as described below (see **Atlases**).

#### Functional image pre-processing

We preprocessed fMRI data using fMRIPrep v25.0.0 (Esteban et al., 2018, 2019, & 2020), which is based on Nipype v1.9.2 (Gorgolewski et al., 2011 & 2018). For each blood-oxygen level dependent (BOLD) run per subject, across all tasks and sessions, the following preprocessing was performed. First, a reference volume was generated for use in head motion correction. Head-motion parameters with respect to the BOLD reference (transformation matrices and six corresponding rotation and translation parameters) are estimated before any spatiotemporal filtering using mcflirt (FSL v6.0.7.7; Jenkinson et al., 2002). The BOLD reference was then co-registered to the T1w reference using bbregister (FreeSurfer), with six degrees of freedom. The aligned T2w image was used for initial co-registration. The BOLD time series were resampled onto the left/right-symmetric template “fsLR” using Connectome Workbench (Glasser et al., 2013). Grayordinates files (Glasser et al., 2013) containing 91k samples were also generated with surface data transformed directly to fsLR 32K space and subcortical data transformed to 2 mm resolution MNI152NLin6Asym space.

#### Functional image post-processing

Following preprocessing with fMRIPrep, fMRI data was post-processed using XCP-D v0.10.7 (Mehta et al., 2024; Ciric et al., 2017; Satterthwaite et al., 2013a), which was built with Nipype v1.10.0 (Gorgolewski et al., 2011 & 2018). Many internal operations of XCP-D use AFNI (Cox, 1996; Cox and Hyde, 1997), Connectome Workbench (Marcus et al., 2011), ANTS (Avants et al., 2008), TemplateFlow v24.2.2 (Ciric et al., 2022), matplotlib v3.10.0 (Hunter, 2007), Nibabel v5.3.2 (Brett et al., 2024), Nilearn v0.11.1 (Abraham et al., 2014), numpy v2.2.1 (Harris et al., 2020), pybids v0.18.1 (Yarkoni et al., 2019), and scipy v1.15.1 (Virtanen et al., 2020).

For each fMRI run that was successfully processed with fMRIPrep, the following post-processing was performed using XCP-D. Non-steady-state volumes were extracted from the preprocessed confounds and were discarded from both the BOLD data and nuisance regressors. The BOLD data were converted to NIfTI format, despiked with AFNI’s 3dDespike, and converted back to CIFTI format. Nuisance regressors were regressed from the BOLD data using a denoising method based on Nilearn’s approach. The time series were band-pass filtered using a second-order Butterworth filter, in order to retain signals between 0.01-0.08 Hz; the same filter was applied to the confounds. The resulting time series were then denoised using linear regression. Task data was post-processed in the same manner as resting-state data without accounting for task-related activation in denoising. Processed functional time series were calculated for parcels in a wide range of atlases (see **Atlases** below) using Connectome Workbench. In cases of partial coverage, uncovered vertices (values of all zeros or NaNs) were either ignored (when the parcel had >50.0% coverage) or were set to zero (when the parcel had <50.0% coverage). Inter-regional functional connectivity between all regions was computed for each atlas as the Pearson correlation between unsmoothed time series of parcel pairs using Connectome Workbench. Denoised BOLD was then smoothed using Connectome Workbench (Marcus et al., 2011) with a Gaussian kernel (full width at half maximum (FWHM) = 6 mm); as noted above, smoothing was applied after parcel time series were extracted for each atlas to avoid blurring across parcel borders.

Additionally, XCP-D outputs include the Amplitude of Low Frequency Fluctuations (ALFF; Zou et al., 2008) and Regional Homogeneity (ReHo; Jiang and Zuo, 2016). ALFF was computed by transforming the mean-centered, standard deviation-normalized, denoised BOLD time series to the frequency domain. The power spectrum was computed within the 0.01-0.08 Hz frequency band and the mean square root of the power spectrum was calculated at each voxel or vertex to estimate ALFF. The resulting ALFF values were then multiplied by the standard deviation of the denoised BOLD time series to retain the original scaling. ALFF maps were smoothed with the Connectome Workbench using a Gaussian kernel (FWHM = 6 mm). In addition, for each hemisphere, ReHo was computed using surface-based 2dReHo (Zhang et al., 2019). Specifically, for each vertex on the surface, Kendall’s coefficient of concordance (KCC) was computed with nearest-neighbor vertices to yield ReHo. For the subcortical, volumetric data, ReHo was computed with neighborhood voxels using AFNI’s 3dReHo (Taylor and Saad, 2013).

#### Task fMRI modeling

We used Nilearn (Nilearn contributors et al., 2025) to fit first-level generalized linear models for fMRI data collected during the fractal *n*-back task (one run per participant). The model included regressors for both the 2-back and 0-back conditions. In the response time-duration model, we included a regressor that models trial onsets with durations equal to participants’ response times (RTDur), following the ConsDurRTDur model described in Mumford et al. (2023). We calculated participant-level effect size maps for the 2-back > 0-back, 2-back > baseline, and 0-back > baseline contrasts. We then used these maps to perform a group-level analysis in Nilearn. Statistical inferences were based on non-parametric permutation testing (10,000 permutations) with voxel family-wise error (FWE) correction (two-sided, p < 0.05, cluster-forming threshold p < 0.001).

We also compared the group-level 2-back > 0-back contrast results from our data with results from a similar fractal *n-*back task administered as part of the Philadelphia Neurodevelopmental Cohort (PNC; Satterthwaite et al., 2016). Both second-level maps (visualized as z-statistics) were defined in the MNI152 space, then resampled to the fsLR surface and parcellated into 400 cortical regions using the Schaefer 400 atlas (Schaefer et al., 2018) with Neuromaps (R. Markello et al., 2024; R. D. Markello et al., 2022). We quantified the spatial correlation with the PNC group-level task contrast map using a Pearson correlation. To test the spatial correspondence of these maps, we used the Alexander-Bloch rotation method (i.e., spin test; Alexander-Bloch et al., 2018). Specifically, we compared the empirical spatial correlation between maps to a null distribution generated using 10,000 spatial autocorrelation-preserving null maps.

#### Diffusion image processing

We used QSIPrep v1.0.0 (Cieslak et al., 2021) based on Nipype v1.9.1 (Gorgolewski et al., 2011 & 2018), to preprocess DWI data. Many internal operations of QSIPrep use Nilearn v0.10.1 (Abraham et al., 2014) and Dipy (Garyfallidis et al., 2014). First, the anatomical reference image was reoriented into AC-PC alignment (an orientation based on the anterior and posterior commissures) via a 6-DOF transform extracted from a full Affine registration to the MNI152NLin2009cAsym template. A full nonlinear registration to the template from AC-PC space was estimated via symmetric nonlinear registration (SyN) using antsRegistration. Brain extraction was performed on the T1w image using SynthStrip (Hoopes et al., 2022) and automated segmentation was performed using SynthSeg (Billot et al., 2023) from FreeSurfer v7.3.1.

DWI data were denoised using the Marchenko-Pastur PCA method implemented in dwidenoise (Tournier et al., 2019; Veraart et al., 2016a; Veraart et al., 2016b; Cordero-Grande et al., 2019) with an automatically-determined window size of 3 voxels. After MP-PCA, Gibbs ringing was removed using TORTOISE (Lee, Novikov, and Fieremans 2021; Irfanoglu et al., 2025). Any images with a *b*-value less than 100 s/mm^2^ were treated as a *b* = 0 image. FSL’s eddy was used for head motion correction and eddy current correction (Andersson and Sotiropoulos 2016). Eddy was configured with a *q*-space smoothing factor of 10, a total of 5 iterations, and 1000 voxels used to estimate hyperparameters. A quadratic first level model and a linear second level model were used to characterize Eddy current-related spatial distortion. Field offset was attempted to be separated from subject movement. *Q*-space coordinates were forcefully assigned to shells; shells were aligned post-eddy. Eddy’s outlier replacement was run (Andersson et al., 2016). As part of outlier replacement, data were grouped by slice, only including values from slices determined to contain at least 250 intracerebral voxels. Groups of voxels deviating by more than 4 standard deviations from the prediction had their data replaced with imputed values. Final interpolation was performed using the Jacobian modulation method.

Framewise displacement (FD) was calculated using the implementation in Nipype (following the definitions by Power et al., 2014). The head-motion estimates calculated in the correction step were also placed within the corresponding confounds file. Slicewise cross correlation was also calculated. The DWI time series were resampled to AC-PC, generating a preprocessed DWI run in AC-PC space with 1.7 mm isotropic voxels. We also calculated Neighboring DWI Correlation (NDC; Yeh et al., 2019), a metric of DWI image quality (Richie-Halford et al., 2022) that measures the mean spatial correlation of a volume between its neighbors in *q*-space.

After preprocessing with QSIPrep, the preprocessed DWI images were reconstructed using multiple complementary methods using QSIRecon v1.1.0 (Cieslak et al., 2021), which is based on Nipype v1.9.1 (Gorgolewski et al., 2011 & 2018). QSIRecon implements multiple models of diffusion data to calculate a rich variety of whole-brain parametric microstructure maps. Brain masks from SynthStrip were used in all subsequent reconstruction steps.

First, matching the workflow from the ABCD study (Hagler et al., 2019; Karcher & Barch, 2021; Volkow et al., 2018), we computed a “full shell” tensor fit (Basser et al., 1994) using data from all *b*-values, using ‘EstimateTensor’ with WLLS regularization. This model is widely used to model Gaussian diffusion data and calculates scalar maps including fractional anisotropy (FA), mean diffusivity (MD), axial diffusivity (AD), and radial diffusivity (RD). These images can be found in the `qsirecon-TORTOISE_model-tensor` directory.

Second, we implemented diffusion kurtosis imaging (DKI: Jensen et al., 2005; Jensen & Helpern, 2010), which fits a fourth-order tensor such that it might be more accurate in cases of non-Gaussian diffusion. DKI fits a model using the full shell with all *b*-values, calculating scalar maps including FA, MD, AD, RD, mean kurtosis (MK), radial kurtosis (RK), and axial kurtosis (AK). The dipy implementation of DKI was used (Henriques et al., 2021) and results can be found in the `qsirecon-DIPYDKI` directory.

Third, we fit the NODDI model (Zhang et al., 2012) using the AMICO implementation (Daducci et al., 2015). A value of 1.7E-03 was used for parallel diffusivity and 3.0E-03 for isotropic diffusivity (Guerrero et al., 2019). Intracellular volume fraction (ICVF), isotropic volume fraction (ISOVF), tissue fraction (1 - ISOVF), modulated ICVF, and Orientation Dispersion (OD) maps were also computed (Parker et al., 2021).

Fourth, we used the Mean Apparent Propagator method (MAPMRI: Özarslan et al., 2013), included in TORTOISE (Irfanoglu et al., 2025). When fitting MAPMRI, we first fit a diffusion tensor using low *b*-value shells to determine the primary coordinate frame of the ensemble average diffusion propagator (EAP). This tensor fit can be considered the “inner shell” complementing the “full shell” tensor fit included in the `qsirecon-TORTOISE_model-tensor` results. Numerous scalar maps can be calculated from a MAPMRI fit, including the following return probabilities: Return to Origin Probability (RTOP), Return to Axis Probability (RTAP), and Return to Plane Probability (RTPP). General Non-Gaussianity (NG), Non-Gaussianity parallel (NGPar), and Non-Gaussianity Perpendicular (NGPer) to the primary direction were calculated. Propagator Anisotropy (PA), and Propagator Anisotropy theta (PAth) were also calculated (Özarslan et al., 2013).

Finally, diffusion orientation distribution functions (ODFs) were reconstructed using generalized q-sampling imaging (GQI: Yeh, Wedeen, and Tseng, 2010) with a ratio of mean diffusion distance of 1.25 in DSI Studio (version 94b9c79). This method also describes more complicated diffusion dynamics without a model. Instead, it uses an analytic transform of the diffusion signal (Yeh, Wedeen, and Tseng., 2010) to calculate water diffusion orientation distribution functions (dODFs), yielding scalar maps such as generalized fractional anisotropy (GFA), quantitative anisotropy (QA), MD, and isotropic component (ISO) (Yeh et al., 2013).

To generate individual white matter bundles, we ran Automatic Tractography in DSI Studio (version 94b9c79) via the AutoTrack implementation. AutoTrack provides bundle shape statistics (Yeh, 2020), such as length, volume, surface area, endpoint surface area, endpoint radius, endpoint irregularity, overall curvature, and overall elongation. The average value of microstructural measures from each reconstruction model described above was also calculated for each bundle. In total, this process yielded 19 macrostructural and 64 microstructural metrics per bundle. We displayed individual white matter bundles that were reconstructed by DSI Studio using a custom Python script implementing features from dipy (Garyfallidis et al., 2014), fury (Garyfallidis et al., 2021), and pyAFQ (Kruper et al., 2021).

#### ASL image processing

We used ASLPrep v0.7.5 (Adebimpe et al., 2022, 2023) based on fMRIPrep (Esteban et al., 2018, 2019, & 2020) and Nipype v1.8.6 (Gorgolewski et al., 2011 & 2018), to process ASL MRI data. Many internal operations of ASLPrep use Nilearn v0.11.1 (Abraham et al., 2014), NumPy (Harris et al., 2020), and SciPy (Virtanen et al., 2020). For each ASL run, the following preprocessing was performed across all tasks and sessions. First, a reference volume was generated to use for head motion correction. Head-motion parameters were estimated for the ASL data using FSL’s mcflirt (Jenkinson et al., 2002). Motion correction was performed separately for tagged and control volumes, in order to account for intensity differences between different contrasts, which can conflate intensity differences with head motion if the contrasts are motion-corrected together (Wang et al., 2008). Next, we concatenated the motion parameters across volume types and re-calculated the relative root mean-squared deviation. The ASL reference was then co-registered to the T1w reference using bbregister (FreeSurfer) with six degrees of freedom. Notably, all resampling in ASLPrep concatenates all transformations so as to only interpolate the data once. Gridded (volumetric) resampling was performed using antsApplyTransforms, configured with Lanczos interpolation to minimize the smoothing effects of other kernels (Lanczos, 1964).

Following preprocessing and as described in detail below, ASLPrep generates cerebral blood flow (CBF) maps using four different methods: standard CBF, SCORE-processed CBF (Structure Correlation based Outlier Rejection CBF), SCRUB CBF (Structural Correlation with RobUst Bayesian CBF), and BASIL CBF (Bayesian Interface for Arterial Spin Labeling CBF). First, standard CBF maps were calculated using a single-compartment general kinetic model (Buxton et al., 1998). Calibration (M0) volumes associated with the ASL scan were smoothed with a Gaussian kernel (FWHM = 5 mm) and the average calibration image was calculated and scaled by 10.0. Second, the Structural Correlation based Outlier Rejection (SCORE) algorithm was applied to the CBF time series to discard CBF volumes with outlying values (Dolui et al., 2017) before computing the mean CBF. Third, we calculated CBF values using both the SCORE and the Structural Correlation with RobUst Bayesian (SCRUB) algorithm. SCRUB uses structural tissue probability maps to reweight the mean CBF (Dolui et al., 2017); note that the CBF time series used to calculate SCRUB CBF was cleaned using SCORE. Finally, CBF was also computed using Bayesian Inference for Arterial Spin Labeling (BASIL; Chappell et al., 2009), as implemented in FSL. BASIL computes CBF using a spatial regularization of the estimated perfusion image and additionally calculates a partial-volume corrected CBF image (Chappell et al., 2011).

For all CBF maps, we calculated the quality evaluation index (QEI; Dolui et al., 2017). QEI scores are based on the similarity between each CBF map and the structural images, the spatial variability of each CBF image, and the percentage of gray matter voxels containing negative CBF values. Parcellated CBF estimates were extracted for multiple atlases (as described below). Uncovered voxels were defined as those with values of all zeros or NaNs. Uncovered voxels were ignored when the parcel had more than 50.0% coverage, or the whole parcel was set to zero when the parcel had less than 50.0% coverage.

#### Atlases

The processed sMRI, fMRI, and ASL data include parcellated data from multiple atlases (**Figure S1**). FreeSurfer data (i.e., sMRI data) was parcellated using various atlases including Desikan Killiany (Desikan et al., 2006), Glasser (Glasser et al., 2016), Gordon (Gordon et al., 2016), Yeo (Yeo et al., 2011), Schaefer 200 through 1000 (Schaefer et al., 2018), and Brodmann areas (Garey, 1994). fMRI data processed with XCP-D and ASL data processed using ASLPrep include parcellated data from many of the same atlases, including Schaefer, Gordon, and Glasser atlases. However, XCP-D and ASLPrep use the Schaefer Supplemented with Subcortical Structures (4S) atlas (Schaefer et al., 2018; Pauli et al., 2018; King et al., 2019; Najdenovska et al., 2018; Glasser et al., 2013) at 10 different resolutions (156, 256, 356, 456, 556, 656, 756, 856, 956, and 1056 parcels) which combines the multi-resolution Schaefer parcellation with 56 subcortical parcels drawn from the CIT168 subcortical atlas (Pauli et al., 2018), Diedrichson cerebellar atlas (King et al., 2019), HCP thalamic atlas (Najdenovska et al., 2018), and the amygdala and hippocampus parcels drawn from the HCP CIFTI subcortical parcellation (Glasser et al., 2013). XCP-D output also contains additional parcellated data from the Masonic Institute for the Developing Brain (MIDB) precision brain atlas (thresholded at 75% probability; Hermosillo et al., 2024), and the Tian subcortical atlas (Tian et al., 2020).

## Data Records

### Neuroimaging Data Structure and Organization

Raw neuroimaging data from this study are provided on OpenNeuro [accession number ds007116; Link]. The original DICOM files were converted to NifTI format and structured to align with the Brain Imaging Data Structure (BIDS, version 1.8.0; Gorgolewski et al., 2016) using HeuDiConv (version 1.3.2; Halchenko et al., 2024). A custom heuristic file (available on this study’s GitHub repository; see **Code Availability**) was generated to specify the organization of files and extract metadata from DICOMs. The extracted metadata was used to generate the JSON sidecars accompanying each NIfTI file in the BIDS dataset. As stipulated by BIDS, files were arranged into subject and session folders, with a `participants.tsv` file detailing demographic information, and an accompanying `sessions.tsv` file specifying relative acquisition times. Acquisition times were anonymized to prevent sharing protected health information such that the first scan date was set to the 15th of the original month in the year 1800. Subsequent sessions were adjusted accordingly such that the relative time intervals between sessions remained accurate, with times rounded to the nearest hour. Detailed information about the files included in the dataset is provided in **Table S2** and **Figure S2.**

In addition to the raw neuroimaging data, we provide analysis-ready processed derivatives on OpenNeuro. Each pipeline produced derivatives in BIDS format, along with HTML summaries of preprocessing steps. The outputs from freesurfer-post are included along with the raw neuroimaging data on OpenNeuro in the `derivatives/freesurfer-post/` folder [ds007116; Link]. The participant-level and group-level task contrast maps from the *n*-back task are also included with the raw neuroimaging data on OpenNeuro in the `derivatives` folder [ds007116; Link]. Group-level task contrast maps are also available on Neurovault [Link]. For the rest of the processing pipelines, imaging derivatives from each BIDS App outputs are uploaded as separate datasets on OpenNeuro, as follows: [sMRIPrep: ds007089, Link]; [fMRIprep: ds007088, Link]; [XCP-D: ds007402, Link]; [QSIPrep: ds007090, Link]; [QSIRecon: ds007091, Link]; [ASLPrep: ds006744, Link].

### Clinical Diagnostic and Phenotypic Data

Information about each participant’s study group (e.g., PS, ADHD, or TD), determined from trained clinical assessors and confirmed by a clinical psychiatrist, is available on OpenNeuro (Markiewicz et al., 2021) [accession number ds007116; Link]. All diagnoses from the clinical interview are specified in the `participants.tsv` file. This file also includes demographic data such as participant age and sex.

Cognitive assessment data from the Penn CNB results and self-report data are available under the `phenotype/` directory of the raw dataset on OpenNeuro [accession number ds007116; Link]. Cognitive assessment data are provided as summary scores based on accuracy and reaction time. Self-report data are provided as raw responses with calculated summary scores for each session. Each Penn CNB task and self-report survey has a separate tsv file and an accompanying JSON file that describes the data represented in each column.

## Technical Validation

### Quality Control

Assessment of data quality is critically important for any neuroimaging study. Imaging artifacts, often due to in-scanner motion, can introduce bias in MRI measures (Elyounssi et al., 2025; Gilmore et al., 2021; Power et al., 2012; Reuter et al., 2015; Rosen et al., 2018; Satterthwaite et al., 2012; Van Dijk et al., 2012; Yan et al., 2013). Populations especially prone to higher in-scanner motion include youth (Afacan et al., 2016; Fair et al., 2013; Satterthwaite et al., 2012; Thomson et al., 2024) and those with psychiatric diagnoses (Makowski et al., 2019) such as ADHD (Pardoe et al., 2016; Shi et al., 2025; Thomson et al., 2024) and psychosis-spectrum disorders (Kong et al., 2014; Pardoe et al., 2016; Yao et al., 2017). Furthermore, discrepancies in QC procedures employed by studies that use the same data can yield divergent results. For this data resource, we calculated multiple indices of image quality and in-scanner motion. Additionally, we provide recommended QC guidelines for each imaging modality, which are publicly shared on GitHub. The total numbers of participants and sessions that passed QC based on our recommendations are summarized in **Figure 3**.

**Figure 3.**
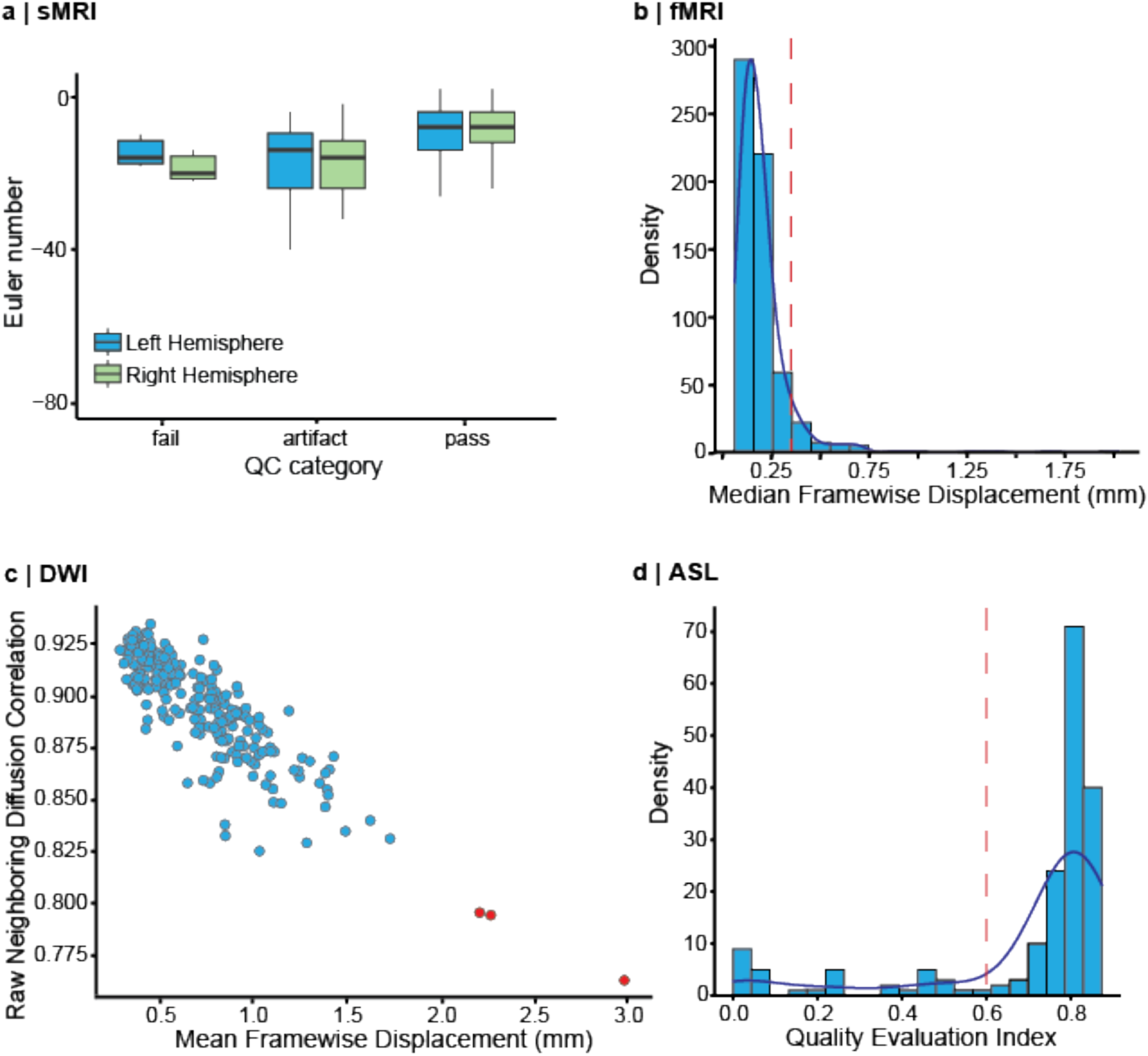
Neuroimaging data quality control. **(a)** Automated image quality measures (i.e., Euler number) along with manual ratings were used to assess the quality of sMRI data. Distribution of left and right hemisphere Euler numbers are visualized using box plots. The Euler number is an automated numerical measure calculated by FreeSurfer and provides a continuous estimate of sMRI data quality. Box plots are grouped according to categorical quality ratings assigned from manual review of T1w scans. Boxes represent the interquartile range of the distribution, where the middle line in each box represents the median Euler number. **(b)** Median FD was used as a measure of fMRI data quality, reflecting participants’ in-scanner motion. Distribution of median framewise displacement (FD) across all functional scans is plotted as a histogram. The red dashed line corresponds to our recommended threshold to consider an fMRI scan of adequate quality (“Pass” label: median FD < 0.3 mm). **(c)** Mean framewise displacement (FD) and raw neighboring DWI correlation (NDC) were used to assess diffusion-weighted imaging (DWI) data quality. Each dot in the scatter plot represents a DWI scan. Red dots indicate ‘outliers’ with low NDC and high motion (i.e., Mean FD > 2.0 mm; NDC < 0.8) that did not “pass” quality control per our recommendations. **(d)** Quality Evaluation Index (QEI) was used as a measure of ASL data quality. Distribution of QEI across all ASL scans is plotted as a histogram. The red dashed line corresponds to our recommended threshold to consider an ASL scan of adequate quality (“Pass” label: QEI > 0.6).

#### sMRI

For T1w structural scans, we provide manual QC ratings from trained raters. Manual ratings were assigned by two reviewers (B.L.S. & T.T.T.), who each rated four slices from each T1w scan in the dataset (two axial slices at z = 20 and z = 37 and two sagittal slices at x = -19 and x = 17) as a binary pass (1) or fail (0) based on visual examination. Following the independent ratings, the two reviewers compared their respective ratings for each individual slice and attempted to reach a consensus. For slices for which there remained a disagreement, a third expert manual rater (M.C.) determined the final rating. Once these final ratings were assigned, we calculated the average rating across all 4 slices for each scan, and used this average to make the final QC recommendation (0 = fail, 1 = pass, between 0 and 1 = artifact). After manual ratings, 86.5% of T1w sessions passed QC (**Figure 4**).

In addition to these manual ratings, we provide left and right hemisphere Euler numbers, an automated numerical measure calculated by FreeSurfer (Dale et al., 1999; Rosen et al., 2018) (**Figure 3a**). The Euler number quantifies the complexity of the surface topology of the reconstructed cortical surface, driven by the number of surface holes present. A value of 2 represents the maximum quality, whereas lower (including negative) values indicate lower quality. This measure has been validated by independent groups as a continuous measure of structural image quality (Elyounssi et al., 2025; Klapwijk et al., 2019; Rosen et al., 2018; Shafiei et al., 2025). As such, the Euler number provides an index of variation in data quality for images that pass manual QC and can be included as a model covariate in analyses.

#### DWI

For DWI scans, we provide QC recommendations based on measures of in-scanner motion and neighboring DWI correlation (NDC). After visual inspection of the scatterplot of mean FD against NDC, we excluded three scans with both high motion and low NDC (**Figure 3c)**. These scans were also excluded from our calculation of group-level scalar maps. After applying these criteria, 98.6% of DWI sessions passed QC (**Figure 4**). In addition to the categorical QC labels, we recommend including FD and/or NDC as covariates in analyses of images that pass QC.

In addition to the global DWI QC, we provide specific QC ratings for white matter bundles reconstructed by tractometry. Specifically, we defined an ‘outlier’ bundle as any bundle whose total volume was greater than or equal to three standard deviations away from the mean total volume for that bundle across subjects and sessions. We summed the number of outliers for each subject and session: a subject/session with greater than 6 outlier bundles was considered an outlier subject/session and received a ‘fail’ rating.

#### fMRI

We used measures of in-scanner motion and field of view coverage for fMRI QC. Specifically, we used median FD (Power et al., 2012) as a summary measure of head movement. After inspecting the distribution of median FD scores across all scans, we assigned ‘pass’ to any scan with median FD < 0.3 mm. In addition, we assessed the percent coverage for each cortical region in the Schaefer 1000 atlas with added subcortical regions (Schaefer et al., 2018). Percent coverage was defined as the proportion of voxels remaining after preprocessing. Regions with <50% coverage were flagged. Subsequently, a scan was assigned a ‘fail’ rating if more than three regions were flagged for insufficient coverage. As such, all fMRI scans that received a final QC determination of ‘pass’ had median FD < 0.3 mm and three or fewer regions with insufficient coverage (**Figure 3b**). Based on this criteria, 94.4% of fMRI sessions passed QC (**Figure 4**). We excluded scans that did not meet this criteria from group-level correlation matrices (excluded runs *n* = 71 across resting and *n*-back scans; **Figure 5**).

**Figure 4.**
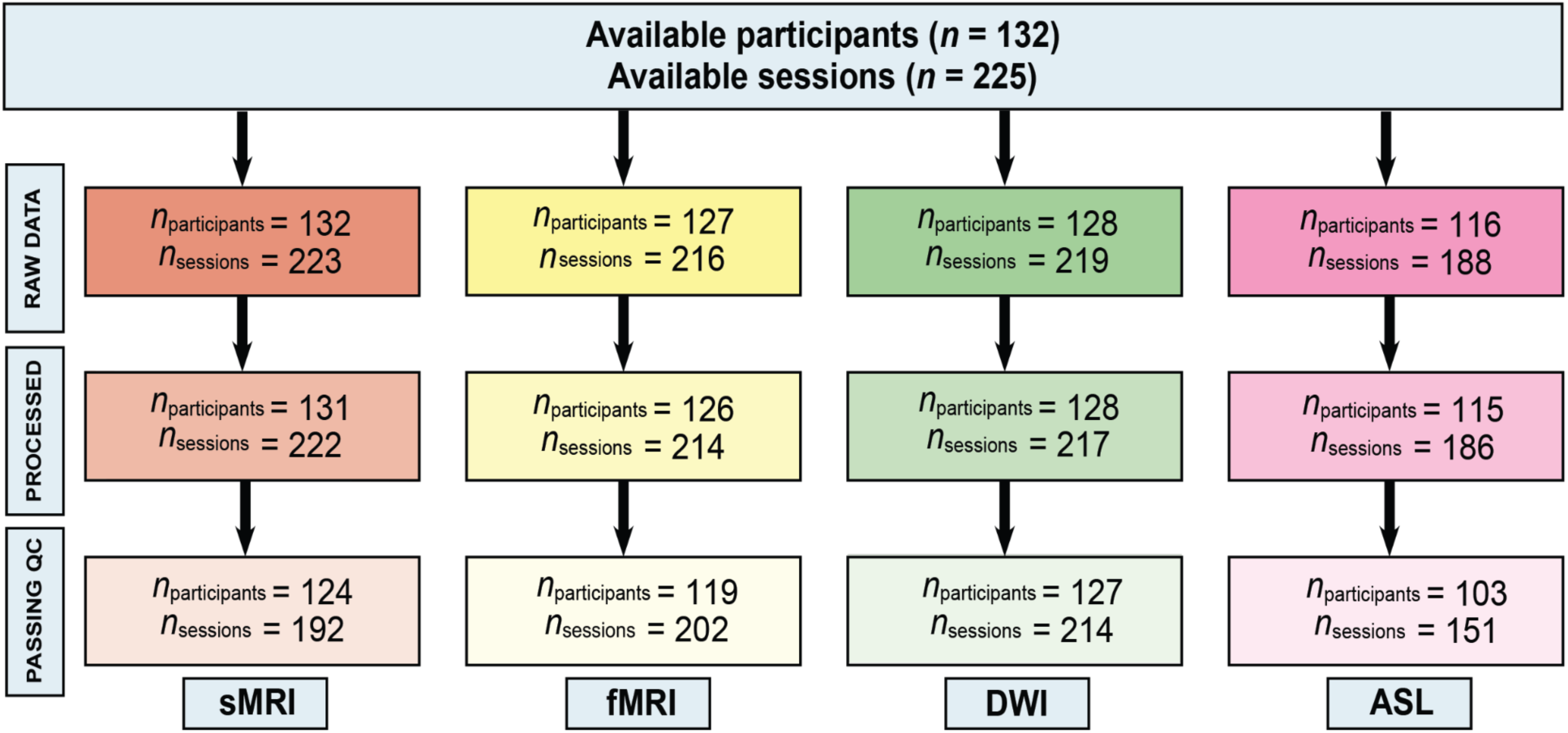
Preprocessing and quality control workflow. The workflow demonstrates the number of participants and scans/sessions from the original collected data that survived preprocessing and met our QC recommendations (i.e., data with “pass” QC).

**Figure 5.**
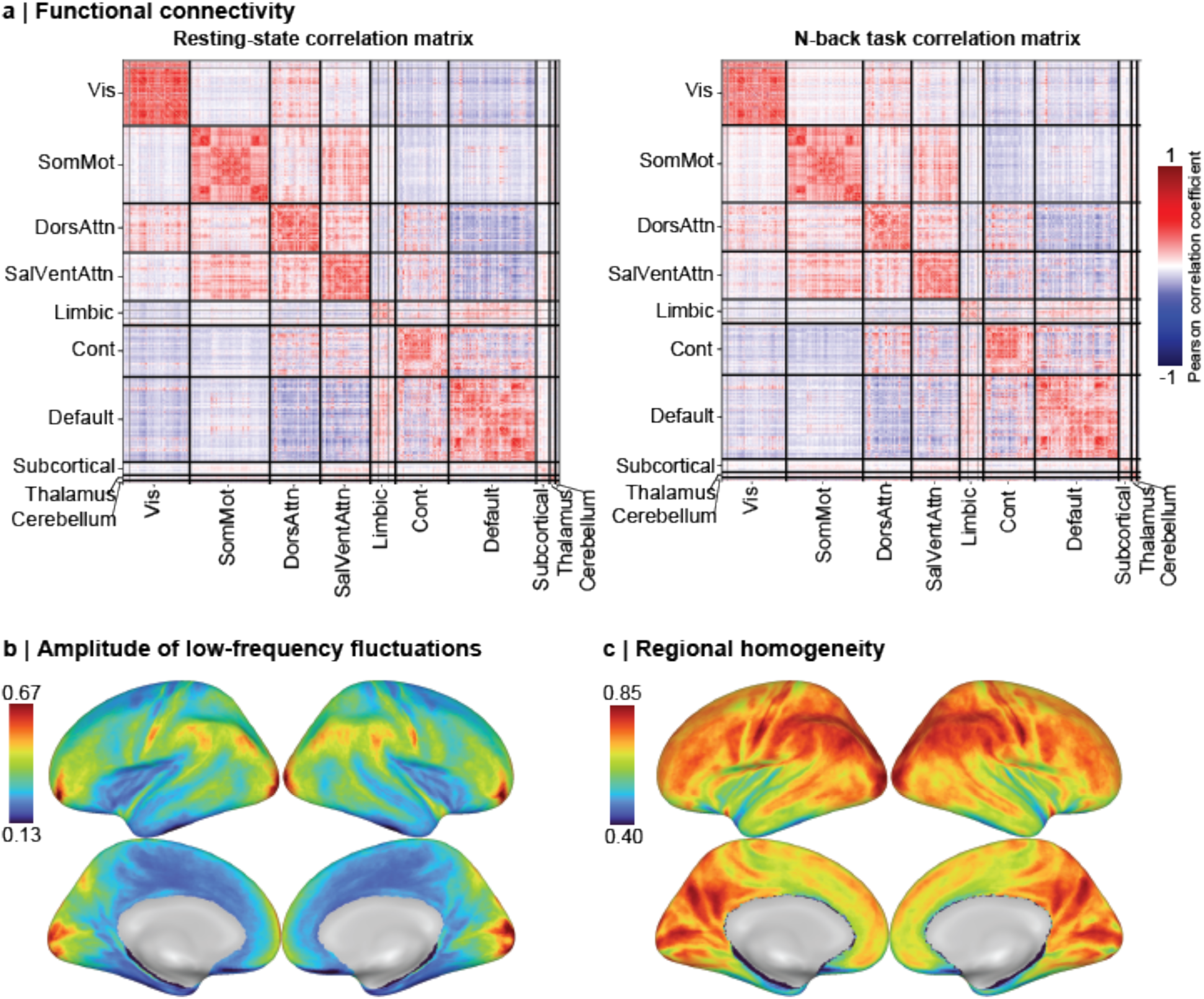
Functional neuroimaging correlation matrices and surface plots. **(a)** Group-level correlation matrices for the resting-state scans and fractal *n*-back task scans, parcellated from the Schaefer 1000 atlas with additional subcortical regions appended and quantified using a Pearson correlation coefficient, revealed canonical functional connectivity patterns and intrinsic networks. Parcels with poor coverage (<50% coverage in more than 3 scans, *n* excluded parcels = 9) across many subjects and sessions were shown as missing values in the group-level correlation matrices. **(b)** Group-level amplitude of low-frequency fluctuations (ALFF) map showed a heterogeneous cortical distribution, with stronger activity in occipital and prefrontal regions. **(c)** Group-level regional homogeneity (ReHo) map showed strong regional coherence of neural activity across the cortex.

#### ASL

For ASL scans, we used QEI (Dolui et al., 2024) as the main QC measure. QEI aggregates various features related to ASL image quality, including spatial similarity of the CBF map to the population-averaged template, the number of voxels with negative CBF values, and the signal-to-noise ratio. QEI values range from 0 to 1, where 1 represents the highest possible quality. Given the distribution of QEI values from our dataset, we assigned a ‘pass’ label to images with a QEI > 0.6 (**Figure 3d**). This threshold was determined based on previous literature classifying an example “average” scan from manual ratings to have a QEI = 0.68 and an example “poor” scan from manual ratings to have a QEI = 0.19 (Dolui et al., 2024). Using this criteria, 82.1% of ASL sessions passed QC (**Figure 4**).

### Neuroimaging Results

#### sMRI

We provide vertex-level sMRI data from FreeSurfer such as cortical thickness, curvature, and sulcal depth, as well as volumetric derivatives such as skull-stripped T1w images, tissue segmentations, brain masks, and the volume of subcortical regions. In addition, freesurfer-post produces atlas-based tabulated measures, including surface area of regions, surface vertices of regions, surface-based total gray matter volume, mean regional cortical thickness and standard deviation, mean curvature, average Gaussian curvature, and regional folding and curvature indices (Van Essen & Drury, 1997). Finally, we include summary whole-brain metrics from multiple atlas parcellations, including the mean and standard deviation of the percentage of all vertices labeled as white and gray matter, ventricle size, cortical thickness, and surface area.

#### fMRI

Functional data derivatives include post-processed clean BOLD time series that are denoised, filtered, and censored; parcellated time series and functional connectivity matrices calculated from Pearson correlations; and ALFF and ReHo derivative maps. We created a group-level correlation matrix, computing the mean for the correlations across all subjects and sessions for both *n*-back and resting-state runs (**Figure 5a**). The mean *n*-back and resting-state correlation matrices revealed canonical functional connectivity patterns, with stronger connectivity within previously described functional communities than between them (Schaefer et al., 2018; Yeo et al., 2011).

In addition to visualizing correlation matrices, we also averaged across every ALFF and ReHo map generated across multiple runs to provide group-level visualizations of ALFF (Zou et al., 2008) and ReHo (Jiang & Zuo, 2016) for resting-state functional MRI data (**Figure 5**). ALFF measures the intensity of BOLD signal fluctuations at each voxel, representing spontaneous neuronal activity at low frequencies. Group-level ALFF map shows a heterogeneous cortical distribution, with higher spontaneous activity in the prefrontal and occipital regions (**Figure 5b**). ReHo measures the local synchronization of a voxel’s time series with its close neighbors. The group-level ReHo map shows generally high values, especially in association cortex (e.g., posterior cingulate and lateral parietal cortex; **Figure 5c**).

Task-contrast derivatives from running first and second-level analyses in Nilearn include both participant-level and group-level contrast maps for the 2-back > 0-back, 2-back > baseline, and 0-back > baseline contrasts. We include maps both with the RTDur regressor (visualized in **Figure 6**), and without the RTDur regressor in our data release. Note that our task design did not allow for robust estimation of baseline. Consistent with this limitation, the 2-back > baseline and 0-back > baseline contrasts showed unexpected deactivations across various modeling strategies (**Figure S3**).

**Figure 6.**
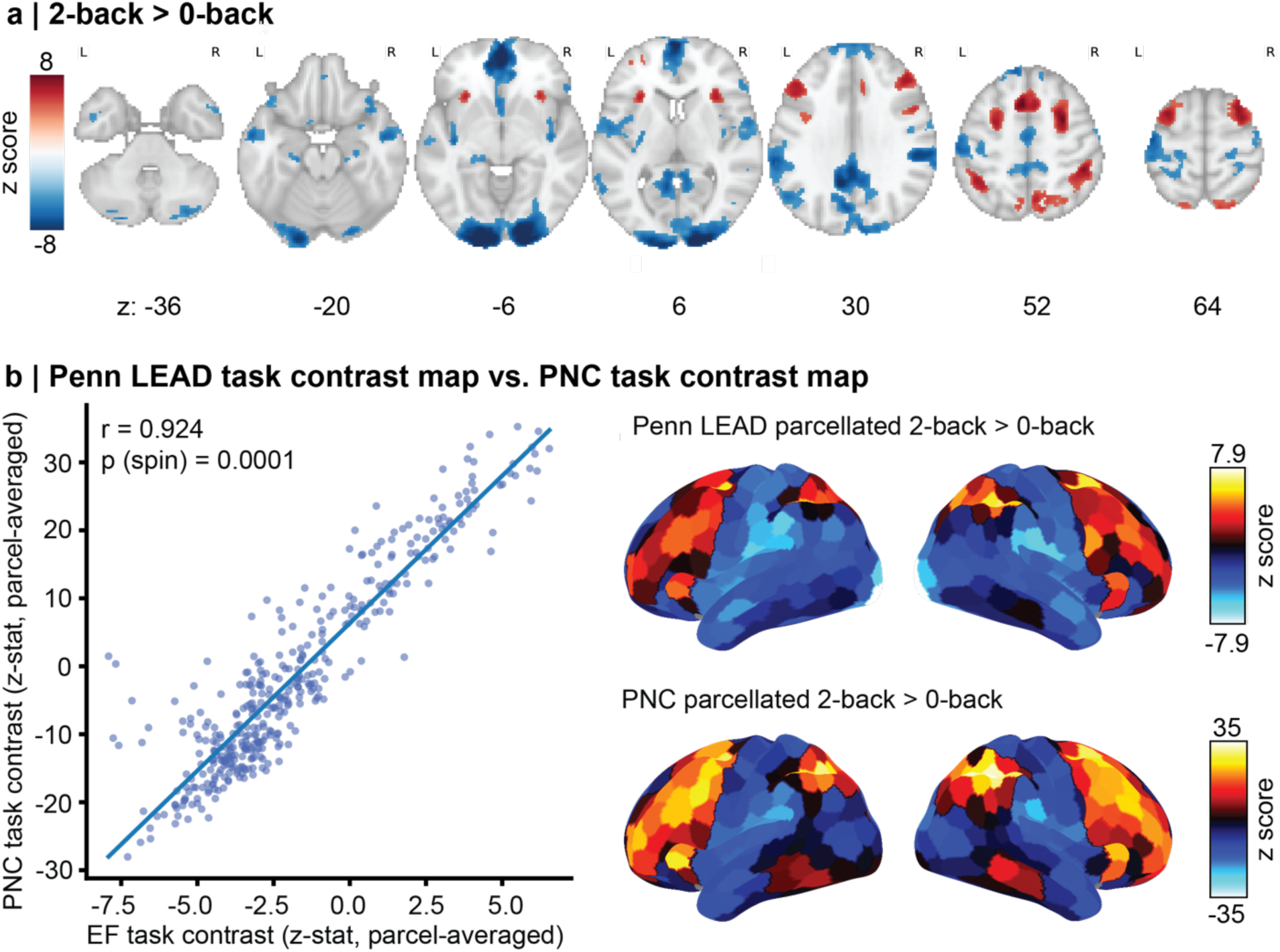
fMRI activation maps from *n*-back task. **(a)** Group-level z-statistic maps for the 2-back > 0-back task contrast from a second-level non-parametric permutation test (10,000 permutations), thresholded at voxelwise FWE-corrected p < 0.05 (two-sided; cluster-forming threshold p < 0.001). **(b)** The relationship between Penn LEAD and Philadelphia Neurodevelopmental Cohort (PNC) z-statistic maps is shown in a scatter plot for the same contrast (left). Both maps were projected to the fsLR surface and parcellated into 400 cortical regions using the Schaefer 400 atlas (right). The statistical significance of the Pearson correlation was assessed using a spin-based permutation test (10,000 permutations). Unthresholded second-level maps were used for the parcellated visualizations and correlation between Penn LEAD and PNC maps.

For the fractal *n-*back working memory task at the group level, the 2-back > 0-back contrast demonstrates increased activation in regions commonly associated with working memory and executive function, including frontoparietal cortical areas (**Figure 6a**), consistent with the extensive prior literature (Huang et al., 2016; Owen et al., 2005). To evaluate the robustness and consistency of these group-level 2-back > 0-back contrast results, we also compared our results to those from a similar fractal *n*-back task administered as part of the Philadelphia Neurodevelopmental Cohort (PNC; Satterthwaite et al., 2016). This revealed that the 2-back > 0-back maps between the two studies were highly similar (**Figure 6b**; *r*=0.924, *p*_spin_=0.0001).

#### DWI

We release derived DWI measures including white matter bundles, bundle-level metrics, and scalar maps from multiple models including traditional tensor-based models, DKI, NODDI, MAPMRI, and GQI. Exemplar group-average maps from one metric per model are displayed in **Figure 7**. From the QSIRecon outputs using the tensor model in DSI Studio, we display the group-level FA map (**Figure 7a**). Similarly, from the GQI model in DSI Studio, we generated group-level GFA maps (**Figure 7b**). As expected, both mean FA and GFA maps showed canonical patterns (Kochunov et al., 2007; Tuch, 2004; Yeh et al., 2010), with maximal values in white matter due to anisotropic diffusion, and lower values in gray matter due to isotropic diffusion. Furthermore, from the DKI model, we generated group-level MK maps that quantify the average degree of non-Gaussian water diffusion, demonstrating expected patterns of higher MK in white matter, and lower values in gray matter (Fieremans et al., 2011; Hansen et al., 2016; Henriques et al., 2021; Jensen et al., 2005; Jensen & Helpern, 2010) (**Figure 7e**). We used NODDI model outputs from QSIRecon (Zhang et al., 2012) to generate group-level ICVF maps, providing an estimate of neurite density within each voxel (**Figure 7c**). The group-level ICVF map demonstrated canonical patterns consistent with previous work (Burzynska et al., 2024; Sadikov et al., 2025; H. Zhang et al., 2012), with higher neurite density in white matter and lower density in gray matter. From the MAPMRI model, we display a group-average RTAP, which quantifies the probability that water molecules return to their original axis (i.e., parallel to the primary diffusion axis). The group-level RTAP map displayed consistent patterns with previous literature (Avram et al., 2022; Özarslan et al., 2013; Sadikov et al., 2025), with higher RTAP values in white matter and lower values in gray matter (**Figure 7d**).

**Figure 7.**
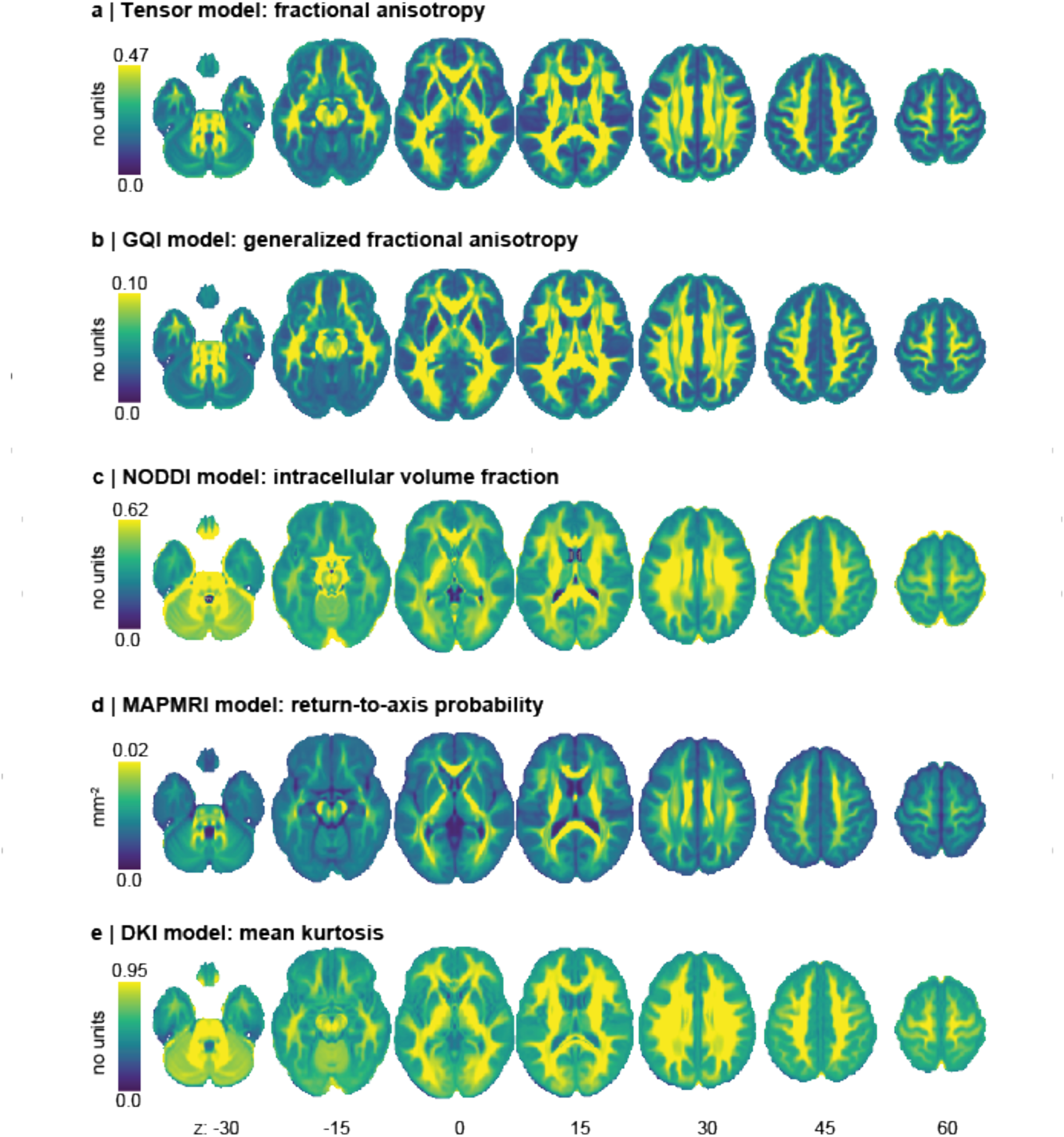
DWI scalar maps. The Penn LEAD dataset provides various DWI scalar maps from QSIRecon. Example group-level maps are shown for four measures: **(a)** Fractional anisotropy from the tensor model in DSI Studio; **(b)** Generalized fractional anisotropy (GFA) from the GQI model in DSI Studio; **(c)** Intracellular volume fraction (ICVF) from the NODDI model; **(d)** Return-to-axis probability (RTAP) from the MAPMRI model with TORTOISE; and **(e)** Mean kurtosis from the DKI model. All axial slices are shown at MNI z-coordinates -30, -15, 0, 15, 30, 45, and 60, overlaid on the MNI152NLin2009cAsym template space.

We also include individual-specific white matter bundles in the data release. In addition to providing the bundles themselves, we release bundle shape statistics as well as the mean value from each scalar metric across all models. Together, this rich data facilitates analysis of person-specific white matter anatomy. Reconstructed white matter bundles (the arcuate fasciculus, corticospinal tract, and inferior fronto-occipital fasciculus) are displayed for two exemplar participants (sub-20188 and sub-20712) across all three longitudinal sessions (**Figure 8**).

**Figure 8.**
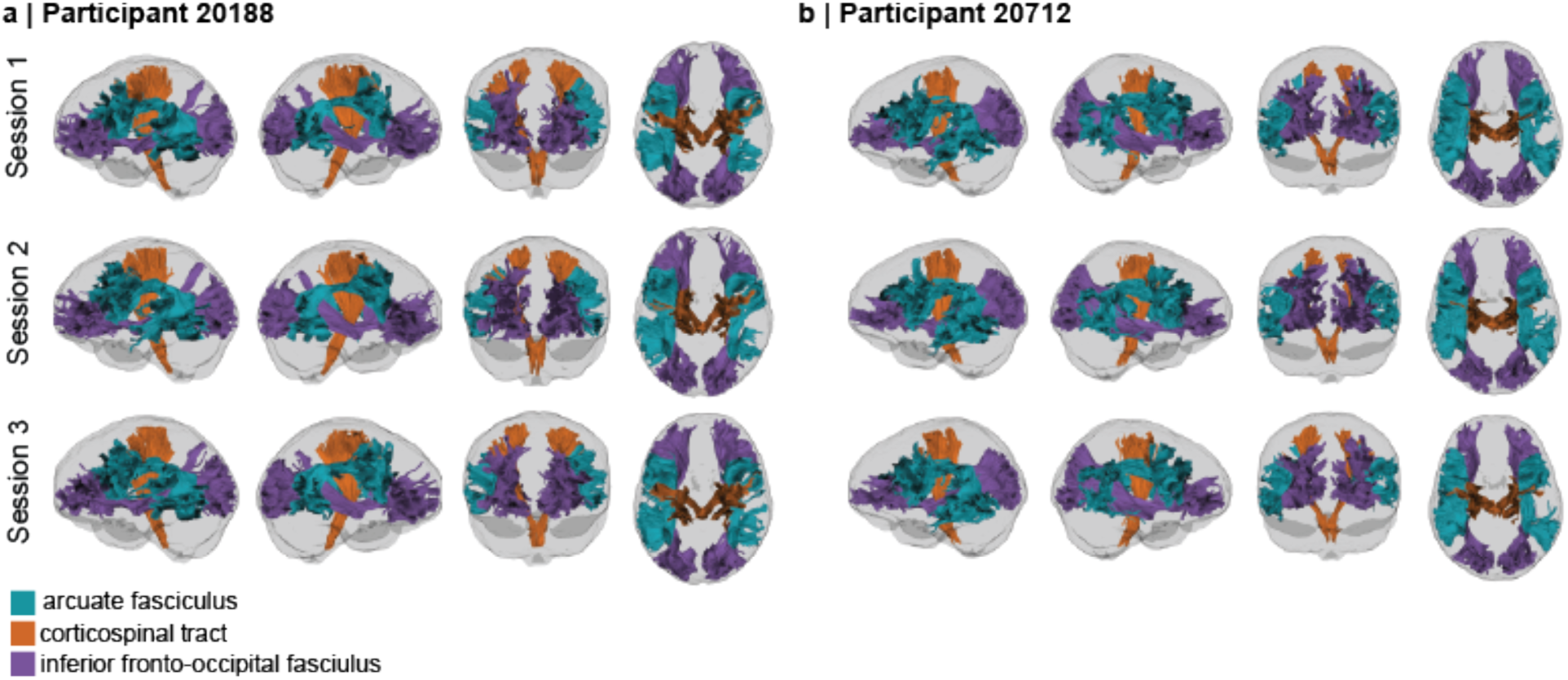
DWI reconstructed tracts. The Penn LEAD dataset also provides individual-level reconstructed diffusion bundles. Reconstructed bundles are shown for two example individuals across all three longitudinal sessions: **(a)** participant 20188 and **(b)** participant 20712. Displayed example bundles are the association arcuate fasciculus (green), projection brainstem corticospinal tract (orange), and association inferior fronto-occipital fasciculus (purple).

#### ASL

ASL derivatives include standard CBF maps, SCORE-processed CBF, SCRUB CBF, BASIL CBF, and BASIL CBF with partial volume correction. The mean CBF maps for some of these methods is displayed in **Figure 9**. CBF maps are consistent with expected patterns of regional cerebral perfusion (Warnert et al., 2020; N. Zhang et al., 2017), with higher CBF in gray matter and lower CBF in white matter.

**Figure 9.**
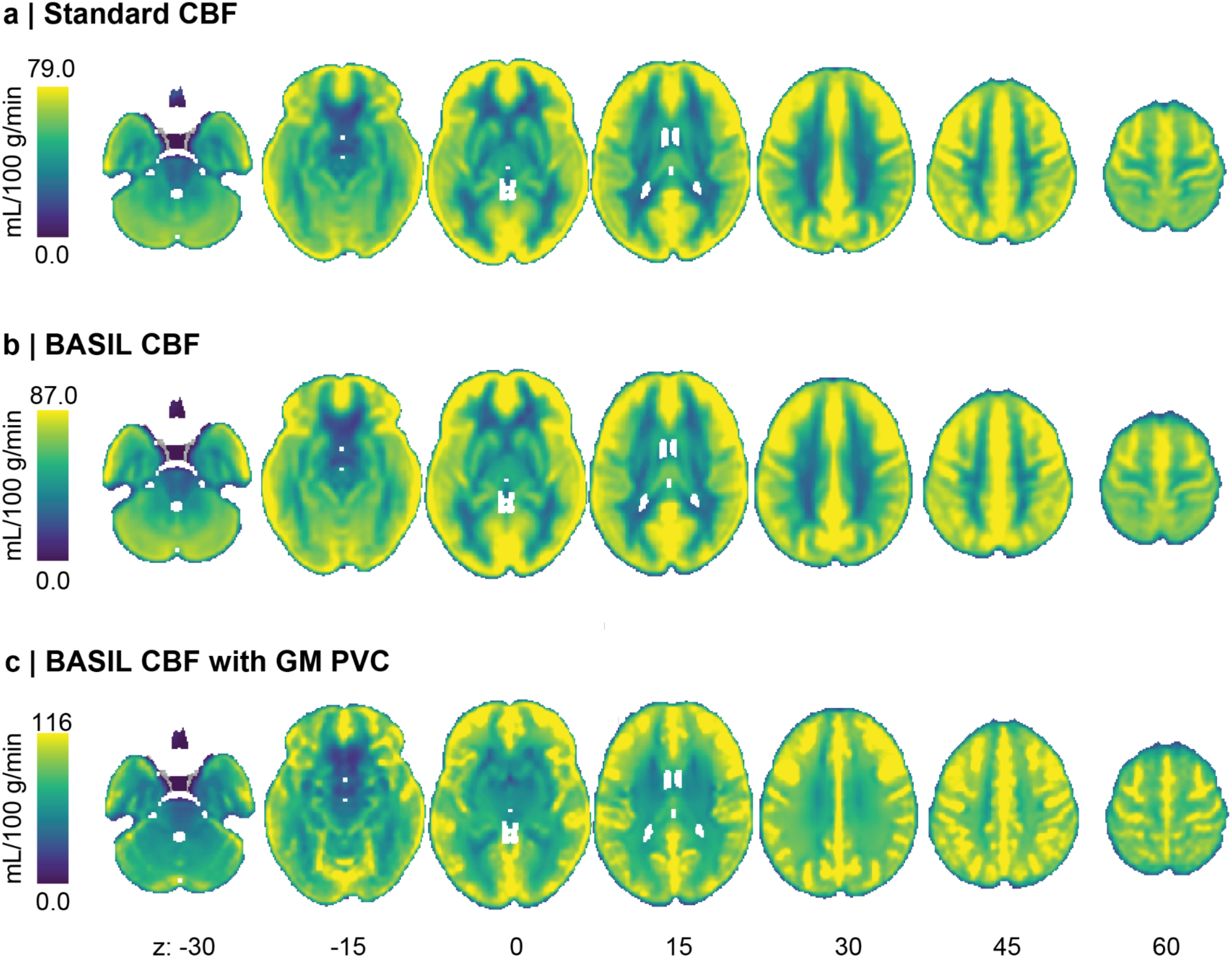
ASL CBF maps. The Penn LEAD dataset provides various measures of cerebral blood flow (CBF) from ASLPrep. Example group-level CBF maps are highlighted for three different measures: **(a)** Standard CBF; **(b)** BASIL estimation of CBF; and **(c)** BASIL CBF estimated with partial volume correction (PVC) for a gray-matter specific map. All axial slices are shown at MNI z-coordinates -30, -15, 0, 15, 30, 45, and 60, overlaid on the MNI152NLin6Asym template.

## Usage Notes

Transdiagnostic cognitive, clinical, and multimodal imaging data are critical for investigating EF deficits in adolescents and understanding how those deficits contribute to diverse psychopathology. The Penn LEAD dataset includes longitudinal curated clinical diagnostic, cognitive, and multimodal neuroimaging data from 132 participants. Participants include adolescents diagnosed with ADHD and PS disorders as well as typically developing individuals. We publicly provide raw, anonymized data in BIDS format for sMRI, fMRI, DWI, ASL, and MEGRE for QSM scans, allowing researchers to process the data through their own pipelines. In addition to the raw data, we provide sMRI, resting-state and task fMRI, DWI, and ASL data derivatives that are fully-processed through BIDS Apps. Processed task fMRI data includes participant and group-level *n*-back task contrast maps. These derived measures – alongside corresponding detailed QC information – ensure that Penn LEAD can immediately be used for hypothesis testing.

## Data Availability

All raw neuroimaging data, behavioral and imaging derivative data from the *n*-back task, phenotypic data, and freesurfer-post outputs are available on OpenNeuro [accession number: ds007116; Link] without a DUA. The raw data also includes output files from the CuBIDS software in the `code/` folder, and the MRI protocol PDF in the `sourcedata/` folder.

Analysis-ready data is also provided on OpenNeuro for sMRIPrep [accession number: ds007089; Link], fMRIPrep [accession number: ds007088; Link], XCP-D [accession number: ds007402; Link], QSIPrep [accession number: ds007090; Link], QSIRecon [accession number: ds007091; Link], and ASLPrep [accession number: ds006744; Link]. Group-level task contrast maps from the *n*-back task are also available on Neurovault [Link], in addition to the derivatives folder of the OpenNeuro dataset above [accession number: ds007116; Link]. Quality control recommendations for each imaging modality are available on GitHub [Link].

## Author Contributions

K.M., S.L., and L.B. contributed to data collection. B.L.S., M.C., S.L.M., T.S., S.P.S., and T.T.T. contributed to data curation. B.L.S., G.S., M.C., S.L.M., T.S., S.P.S., and T.T.T. contributed to analysis. T.D.S. and D.B. acquired funding. T.D.S. provided supervision. B.L.S., G.S., S.P.S., and T.S. contributed to visualization. B.L.S, G.S., and T.D.S. wrote the original draft, and all authors contributed to review and editing of the final draft.

## Code Availability

We used open-source, containerized, reproducible processing pipelines and software for all imaging processing and analysis. HeuDiConv was used to convert images from DICOM to BIDS format [code here; documentation here] (Halchenko et al., 2024). CuBIDS software was used to facilitate curation of BIDS images [code here; documentation here] (Covitz et al., 2022). BIDS-App Bootstrap (BABS) software was used to preprocess images by wrapping around BIDS Apps [code here; documentation here] (Zhao et al., 2024). The custom configuration YAML files used with BABS can be found on GitHub here.

The following BIDS Apps were used: fMRIPrep [code here; documentation here; Docker Hub link here] (Esteban et al., 2018, 2019, 2020), XCP-D [code here; documentation here; Docker Hub link here] (Mehta et al., 2023), QSIPrep [code here; documentation: here; Docker Hub link here] (Cieslak et al., 2021), QSIRecon [code here; documentation here; Docker Hub link here] (Cieslak et al., 2021), freesurfer-post [code here; Docker Hub link here], and ASLPrep [code here; documentation here; Docker Hub link here] (Adebimpe et al., 2022, 2023).

For task contrast analyses, we used Nilearn [code here; documentation here]. All custom code for data organization, quality control, self-report scoring, group-level analyses, and figure generation is available and described in detail on GitHub here.

