## Supplementary Figures and Tables for "A longitudinal data resource to study brain development and transdiagnostic variation in executive function"

**Supplementary Information**

| Image Type | Number of scans in total | Percentage of scans in dominant group |
|---|---|---|
| T1w (normalized) | 223 | 99.1% |
| T2w (normalized) | 224 | 100% |
| T2w (un-normalized) | 220 | 100% |
| Diffusion | 219 | 89.95% |
| Resting state fMRI run 1 | 206 | 87.86% |
| Resting state fMRI run 2 | 199 | 84.42% |
| Resting state fMRI run 3 | 9 | 77.78% |
| Task fMRI run 1 | 209 | 88.52% |
| ASL | 188 | 60.11% |
| m0 scan | 188 | 60.11% |

**Table S1. CuBIDS summary table**. Percentage of sessions belonging to the 'dominant' group for each image modality.

| Data Files | Description |
|---|---|
| participants.tsv | Participant demographics including participant ID, study group, sex, race, ethnicity, and age at intake, as well as clinical diagnostic information |
| participants.json | Describes columns in participants.tsv file |
| dataset_description.json | Describes metadata for entire dataset, including acknowledgements, the version of bids used, funding, and licensing |
| sub-*/sub-*_sessions.tsv | Describes session ID and anonymized acquisition time for each session for that subject |
| sub-*/ses-*/anat/sub-*_ses-*_rec-norm_run-*_T1w.[nii.gz\|json] | Intensity-normalized T1w anatomical NIfTI scan and JSON metadata |
| sub-*/ses-*/anat/sub-*_ses-*_rec-norm_run-*_T2w.[nii.gz\|json] | Intensity-normalized T2w anatomical NIfTI scan and JSON metadata |
| sub-*/ses-*/anat/sub-*_ses-*_run-*_T2w.[nii.gz\|json] | Un-normalized T2w anatomical NIfTI scan and JSON metadata |
| sub-*/ses-*/dwi/sub-*_ses-*_run-*_dwi.bval | DWI b-Values |
| sub-*/ses-*/dwi/sub-*_ses-*_run-*_dwi.bvec | DWI b-Vector |
| sub-*/ses-*/dwi/sub-*_ses-*_run-*_dwi.[nii.gz\|json] | DWI NIfTI scan and JSON metadata |
| sub-*/ses-*/fmap/sub-*_ses-*_acq-dwi_dir-AP_run-*_epi.[nii.gz\|json] | EPI field map for DWI scans with phase encoding in the anterior-posterior direction and JSON metadata |
| sub-*/ses-*/fmap/sub-*_ses-*_acq-dwi_dir-PA_run-*_epi.[nii.gz\|json] | EPI field map for DWI scans with phase encoding in the posterior-anterior direction and JSON metadata |
| sub-*/ses-*/fmap/sub-*_ses-*_acq-fmri_dir-AP_run-*_epi.[nii.gz\|json] | EPI field map for fMRI scans with phase encoding in the anterior-posterior direction and JSON metadata |
| sub-*/ses-*/fmap/sub-*_ses-*_acq-fmri_dir-PA_run-*_epi.[nii.gz\|json] | EPI field map for fMRI scans with phase encoding in the posterior-anterior direction and JSON metadata |
| sub-*/ses-*/func/sub-*_ses-*_task-nback_run-*_bold.[nii.gz\|json] | *n*-back fMRI NIfTI scan and JSON metadata |
| sub-*/ses-*/func/sub-*_ses-*_task-rest_run-*_bold.[nii.gz\|json] | resting-state fMRI NIfTI scan and JSON metadata |

| | |
|---|---|
| sub-*/ses-*/func/sub-*_ses-_task-nback_run*_events.[tsv\|json] | Description of fractal *n*-back task event parameters including stimulus onset, duration, trial type, results, response time, and score. |
| sub-*/ses-*/perf/sub-*_ses-*_run-*_asl.[nii.gz\|json] | ASL NIfTI scan and JSON metadata |
| sub-*/ses-*/perf/sub-*_ses-*_run-*_m0scan.[nii.gz\|json] | reference m0 NIfTI scan and JSON metadata |
| sub-*/ses-*/anat/sub-*_ses-*_echo-*_part-*_MEGRE.[nii.gz\|json] | MEGRE sequence for QSM NIfTI scan and JSON metadata. Part-phase is the phase image, and part-mag is the magnitude image. |

**Table S2. Description of files in BIDS dataset.**

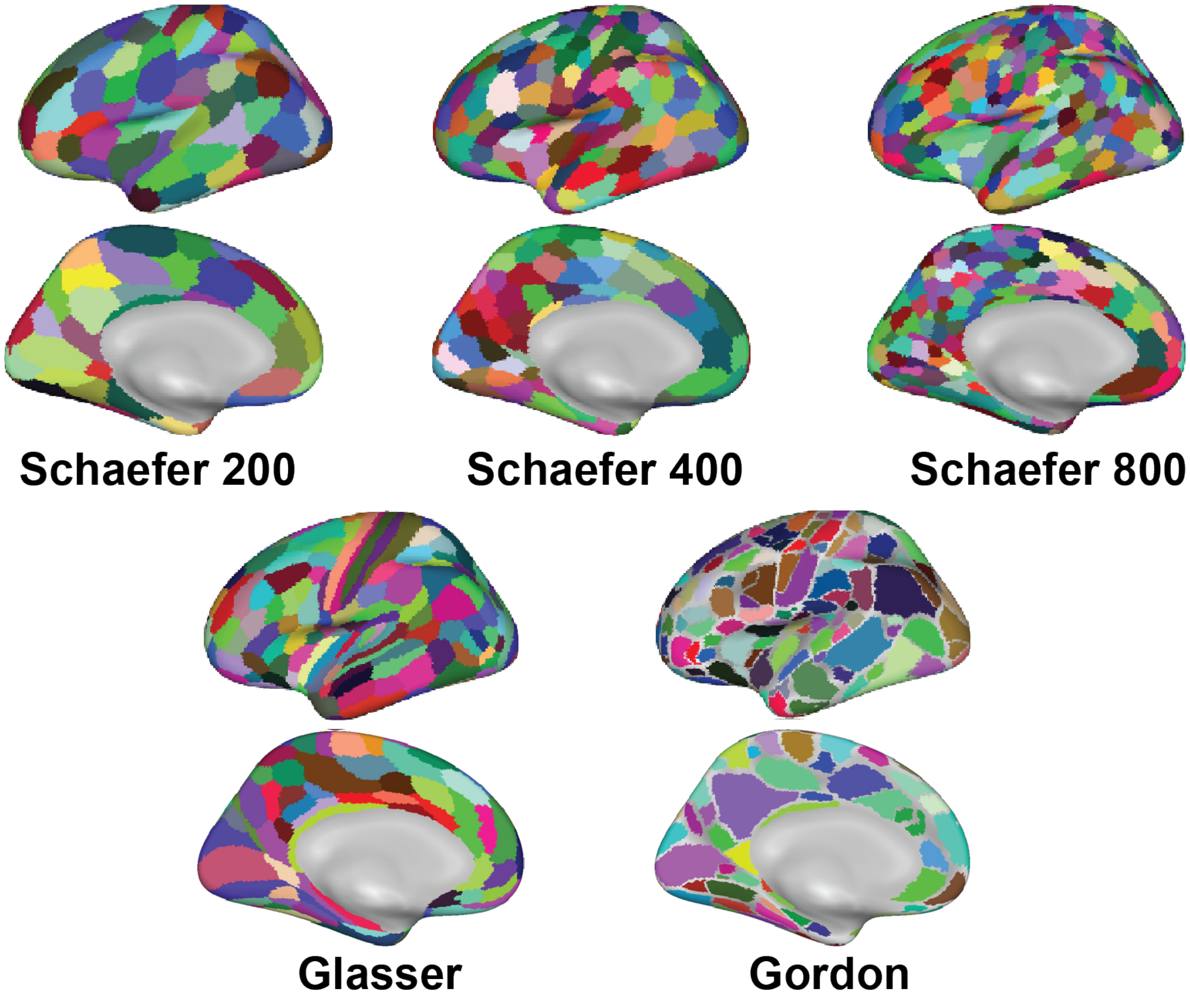

**Figure S1. Example atlases included in the Penn LEAD dataset.** A few examples of atlases used to create parcellated data in the Penn LEAD dataset are highlighted. These atlases include multiple iterations of the Schaefer atlas 100 to 1000 (200, 400, and 800 shown; Schaefer et al., 2018) with subcortical regions appended, the Glasser atlas (Glasser et al., 2016), and the Gordon atlas (Gordon et al., 2016). Additional atlases (not shown) include the MIDB (Hermosillo et al., 2024), HCP (Glasser et al., 2013), and Tian atlases (Tian et al., 2020).

```
├── anat
│   ├── sub-20350_ses-1_echo-1_part-mag_MEGRE.json
│   ├── sub-20350_ses-1_echo-1_part-mag_MEGRE.nii.gz
│   ├── sub-20350_ses-1_echo-1_part-phase_MEGRE.json
│   ├── sub-20350_ses-1_echo-1_part-phase_MEGRE.nii.gz
│   ├── sub-20350_ses-1_echo-2_part-mag_MEGRE.json
│   ├── sub-20350_ses-1_echo-2_part-mag_MEGRE.nii.gz
│   ├── sub-20350_ses-1_echo-2_part-phase_MEGRE.json
│   ├── sub-20350_ses-1_echo-2_part-phase_MEGRE.nii.gz
│   ├── sub-20350_ses-1_echo-3_part-mag_MEGRE.json
│   ├── sub-20350_ses-1_echo-3_part-mag_MEGRE.nii.gz
│   ├── sub-20350_ses-1_echo-3_part-phase_MEGRE.json
│   ├── sub-20350_ses-1_echo-3_part-phase_MEGRE.nii.gz
│   ├── sub-20350_ses-1_echo-4_part-mag_MEGRE.json
│   ├── sub-20350_ses-1_echo-4_part-mag_MEGRE.nii.gz
│   ├── sub-20350_ses-1_echo-4_part-phase_MEGRE.json
│   ├── sub-20350_ses-1_echo-4_part-phase_MEGRE.nii.gz
│   ├── sub-20350_ses-1_rec-defaced_run-01_T2w.json
│   ├── sub-20350_ses-1_rec-defaced_run-01_T2w.nii.gz
│   ├── sub-20350_ses-1_rec-normdefaced_run-01_T2w.json
│   ├── sub-20350_ses-1_rec-normdefaced_run-01_T2w.nii.gz
│   ├── sub-20350_ses-1_rec-normrefaced_run-01_T1w.json
│   └── sub-20350_ses-1_rec-normrefaced_run-01_T1w.nii.gz
├── dwi
│   ├── sub-20350_ses-1_run-01_dwi.bval
│   ├── sub-20350_ses-1_run-01_dwi.bvec
│   ├── sub-20350_ses-1_run-01_dwi.json
│   └── sub-20350_ses-1_run-01_dwi.nii.gz
├── fmap
│   ├── sub-20350_ses-1_acq-dwi_dir-AP_run-01_epi.json
│   ├── sub-20350_ses-1_acq-dwi_dir-AP_run-01_epi.nii.gz
│   ├── sub-20350_ses-1_acq-dwi_dir-PA_run-01_epi.json
│   ├── sub-20350_ses-1_acq-dwi_dir-PA_run-01_epi.nii.gz
│   ├── sub-20350_ses-1_acq-fmri_dir-AP_run-01_epi.json
│   ├── sub-20350_ses-1_acq-fmri_dir-AP_run-01_epi.nii.gz
│   ├── sub-20350_ses-1_acq-fmri_dir-PA_run-01_epi.json
│   └── sub-20350_ses-1_acq-fmri_dir-PA_run-01_epi.nii.gz
├── func
│   ├── sub-20350_ses-1_task-nback_run-01_bold.json
│   ├── sub-20350_ses-1_task-nback_run-01_bold.nii.gz
│   ├── sub-20350_ses-1_task-nback_run-01_events.json
│   ├── sub-20350_ses-1_task-nback_run-01_events.tsv
│   ├── sub-20350_ses-1_task-rest_run-01_bold.json
│   ├── sub-20350_ses-1_task-rest_run-01_bold.nii.gz
│   ├── sub-20350_ses-1_task-rest_run-02_bold.json
│   └── sub-20350_ses-1_task-rest_run-02_bold.nii.gz
└── perf
    ├── sub-20350_ses-1_run-01_aslcontext.tsv
    ├── sub-20350_ses-1_run-01_asl.json
    ├── sub-20350_ses-1_run-01_asl.nii.gz
    ├── sub-20350_ses-1_run-01_m0scan.json
    └── sub-20350_ses-1_run-01_m0scan.nii.gz
```

**Figure S2. Exemplar subject directory structure.** The figure above shows an example folder structure in BIDS format for one session in one subject (sub-20350, ses-1).

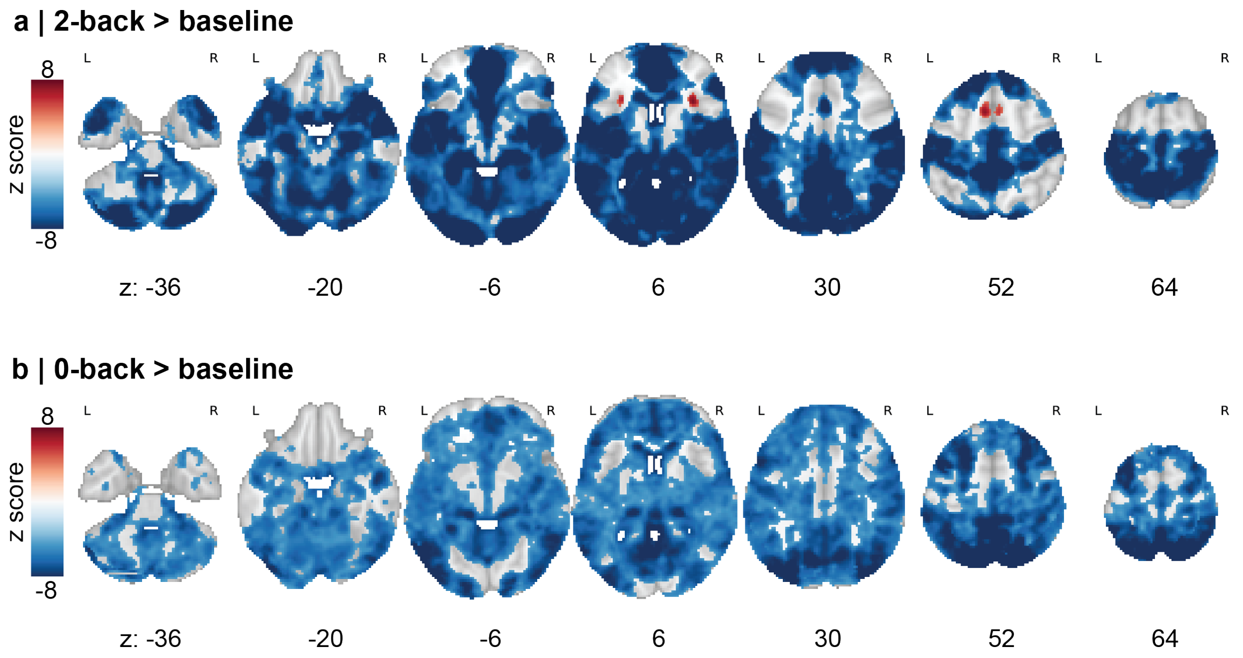

**Figure S3. fMRI activation maps from *n*-back task.** Maps are masked using a voxelwise family-wise error (FWE) corrected $p < 0.05$ (two-sided), with a cluster-forming threshold of $p < 0.001$ based on permutation-derived null distribution. Note that these baseline contrasts are not interpretable, as our task design did not allow for a robust estimation of baseline. **(a)** Group-level z statistic activation maps in MNI152 space for 2-back > baseline task contrast derived from second-level non-parametric permutation testing (10,000 permutations). **(b)** Group-level z statistic activation maps in MNI152 space for 0-back > baseline task contrast derived from second-level non-parametric permutation testing (10,000 permutations).
